# Metamorphosis of memory circuits in *Drosophila* reveal a strategy for evolving a larval brain

**DOI:** 10.1101/2022.06.09.495452

**Authors:** James W. Truman, Jacquelyn Price, Rosa L. Miyares, Tzumin Lee

**Affiliations:** Janelia Research Campus, H.H.M.I., Ashburn, VA; Friday Harbor Laboratories, University of Washington, Friday Harbor, WA 98250

## Abstract

Insects like *Drosophila* produce a second brain adapted to the form and behavior of a larva. Neurons for both larval and adult brains are produced by the same stem cells (neuroblasts) but the larva possesses only the earliest born neurons produced from each. To understand how a functional larval brain is made from this reduced set of neurons, we examined the origins and metamorphic fates of the neurons of the larval and adult mushroom body circuits. The adult mushroom body core is built sequentially of γ Kenyon cells, that form a medial lobe, followed by α’β’, and αβ Kenyon cells that form additional medial lobes and two vertical lobes. Extrinsic input (MBINs) and output (MBONs) neurons divide this core into computational compartments. The larval mushroom body contains only γ neurons. Its medial lobe compartments are roughly homologous to those of the adult and same MBONs are used for both. The larval vertical lobe, however, is an analogous “facsimile” that uses a larval-specific branch on the γ neurons to make up for the missing α’β’, and αβ neurons. The extrinsic cells for the facsimile are early-born neurons that trans-differentiate to serve a mushroom body function in the larva and then shift to other brain circuits in the adult. These findings are discussed in the context of the evolution of a larval brain in insects with complete metamorphosis.

## Introduction

*Drosophila*, like other insects with complete metamorphosis, makes two different brains during its lifetime: one that functions during its larval stage and the other in the adult. Such holometabolous insects evolved from direct developing ancestors (hemimetabolous), which produce only a single brain that functions both in the growing nymphal stages and in the reproductive adult. Even circuits involved in adult-specific behaviors such as flight and oviposition are in place at hatching in these insects and their component neurons change little, except for size from hatching to adulthood (reviewed in Truman, 2005). How then did this single brain arrangement evolve into a two-brain system seen in *Drosophila*? The central brain of the fly larva has only about ten percent of the neurons found in the adult. This numerical difference results from a global arrest of neurogenesis that occurs during embryogenesis, thereby allowing a much shorter time to hatching as compared to direct developing insects but resulting in a nervous system with vastly fewer neurons than found in a hemimetabolous nymph.

While producing such a “mini-brain” is in line with a simplified larval body plan, the truncation of neurogenesis poses challenges for generating the diversity of neuron types needed to make a CNS. It was shown almost 40 years ago (Thomas *et al.,* 1984) that insects have an ancient and highly conserved ground plan for making their nervous system. Neuronal cell types are generated in a modular fashion, with each neural stem cell, called a neuroblast, generating a characteristic lineage of neuron types, with no crosstalk between lineage modules (Taghert & Goodman, 1984). Neuronal identity is established within each lineage through a common temporal code, based on relative birth order (Doe, 2017; Miyares & Lee, 2019; Rossi *et al*., 2021). The evolution of the larval nervous system then occurred within the context of these spatial and temporal mechanisms.

The arrays of neuroblasts in both the central brain (Urbach & Technau, 2003) and ventral nerve cord (VNC) (Thomas *et al.,* 1984; Truman & Ball, 1998) are highly conserved throughout the insects. For the earliest-born neurons, the relative timing of when different cell types arise within a lineage is similar in both flies and direct-developing insects like grasshoppers (*e.g.,* Thomas *et al.,*1984; Jacobs & Goodman, 1989). Consequently, an early truncation of neurogenesis leaves the larva with only early-born cell types with which to make its nervous system.

The problem, then, is how does the larva manage to make a complex brain from this reduced number of cell types? One strategy would be to evolve a new set of temporal rules for making the larval neurons and then “resetting” the system at hatching and making new neurons for the adult CNS. However, an analysis of the genes involved in temporal identity do not show a reset at hatching (Tsuji *et al*., 2008), and, while some larval neurons do die at metamorphosis, most larval neurons, especially those in the brain, persist through metamorphosis and are used again in the adult (Truman, 2005; Roy *et al*., 2007). Some of these neurons have similar functions in both larva and adult, as shown by the demonstration that interneurons causing backward crawling in the larva also control backward walking in the adult (Lee & Doe, 2021). However, as evident in the present study, other neurons have profoundly different functions in the two life stages.

While there have been numerous studies describing how individual neurons change through metamorphosis (e.g., Truman & Reiss, 1976; Levine & Truman, 1985; Roy *et al.,* 2007), they do not reveal why some neurons maintain their function while other radically change as they progress from larva to adult. We decided to examine the metamorphic fates of the assemblage of neurons that provide the input and output for the mushroom bodies. This brain region is specialized in both larva and adult to associate odors with either rewards or punishments and to adjust the animal’s future behavior accordingly (Cognigni, *et al.,* 2018; Thum & Gerber, 2019). Moreover, the wiring diagram of the mushroom body circuitry is known at the EM level for both the larva [Eichler *et al.,* 2017] and adult [Zheng *et al.,* 2017; Li *et al.,* 2020]. In direct developing insects like crickets, the mushroom bodies are assembled during embryogenesis, with different neuron types being produced as embryogenesis progresses (Malaterre *et al.,* 2002). However, the early arrest of neurogenesis in *Drosophila*, results in only the earliest of these cell types being available for making the larval mushroom bodies. We find that this temporal constraint underlies the diversity of functional changes that are seen during metamorphosis.

## Results

### The structure and metamorphosis of the Mushroom Bodies

The core of the *Drosophila* mushroom body is a set of hundreds (larva) to thousands (adult) of small neurons called Kenyon cells (Fig 1A). Their dendrites form the calyx neuropil, which receives olfactory input from the antennal lobes, and their bundled axons extend down the peduncle and into the vertical and medial lobes. The larva has only γ Kenyon cells, which are born during embryogenesis and the first half of larval life. These neurons have bifurcated axons that form the larval vertical and medial lobes. Early in the last larval stage, the mushroom body neuroblasts switch to making α’β’ Kenyon cells and then, at the start of metamorphosis, to αβ Kenyon cells (Lee *et al.,* 1999). The latter two types of neurons also have bifurcated axons with vertical and medial branches, but they remain immature until metamorphosis. At the start of metamorphosis, the γ neurons prune back their axon branches and then regrow only the adult medial branch, while the α’β’ and αβ neurons undergo their maturation (Lee *et al.,* 1999; Awasaki & Ito, 2004). Consequently, the adult mushroom body has three major classes of Kenyon cells, the γ, the α’β’, and the αβ neurons, whose axons form three medial lobes (γ, β’, β) and two vertical lobes (α, α’).

**Figure 1.**
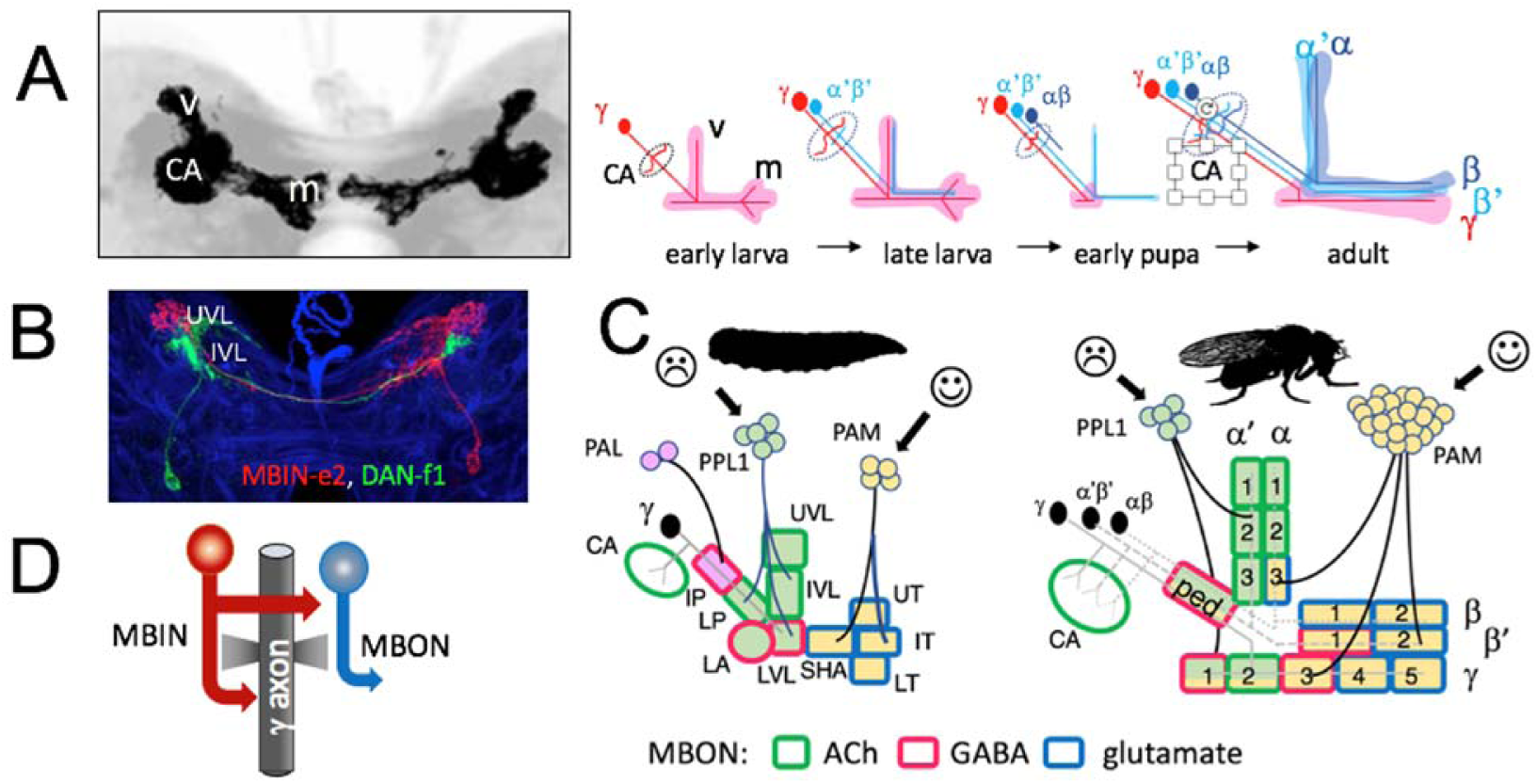
The organization of the larval and adult mushroom bodies in the *Drosophila* brain. **A.** Structure and development of the intrinsic cells (Kenyon cells) of the mushroom body through larval life and metamorphosis. The confocal projection shows a larval brain with the paired mushroom bodies; dendrites project to the calyx (CA) and the axons bifurcate into the vertical (v) and medial (m) lobes. Schematic shows the anatomy of the γ Kenyon cells found in the larva; these are later joined by α’β’ and αβ Kenyon cells to form the medial (γ, β’, β) and vertical (α’, α) lobe systems of the adult. **B.** Projection of a multicolor flip-out (MCFO) image from larval brain showing two mushroom body input neurons that project bilaterally to the upper (UVL) and intermediate (IVL) compartments of the vertical lobes. **C.** The mushroom body peduncle (ped) and lobe systems are divided into computational compartments defined by the projections from three clusters of aminergic neurons, the PAL (light red), PPL1 (light green) and PAM (yellow) clusters. PPL1 input largely indicates punishment, and PAM input indicates reward. These interact with Kenyon cell axons and mushroom body output neurons (MBONs) to cause either avoidance or attraction. The MBON transmitter is indicated by the outline of each compartment. The adult has 16 compartments compared to the ten compartments of the larva. Larval compartments: CA: calyx; IP and LP: intermediate and lower peduncle; LA: lateral appendix; UVL, IVL, LVL: upper, intermediate and lower vertical lobe; SHA: shaft; UT, IT, LT: upper, intermediate and lower toe. **D.** Schematic of the microcircuitry of larval and adult compartments.

The mushroom bodies mediate olfactory learning *via* the set of extrinsic input and output neurons that innervate the peduncle and lobes (Fig 1B). In both larvae and adults, these input and output neurons divide the lobes into non-overlapping, functional compartments, that have a common microcircuit motif (Fig 1C,D) (Eichler *et al.,* 2017; Zheng *et al.,* 2018). Each compartment is defined by the axonal tuft of an aminergic input cell which synapses onto Kenyon cell axons and onto a dedicated output neuron(s). The Kenyon cell axons synapse onto each other, onto the output cell and back onto the input cells. The majority of the <u>m</u>ushroom <u>b</u>ody input <u>n</u>eurons (MBINs) are <u>d</u>op<u>a</u>minergic <u>n</u>eurons (DANs) but a few are <u>o</u>ctop<u>a</u>minergic <u>n</u>eurons (OANs). The DANs come from two clusters, one that generally encodes reward (the PAM cluster) and the other that mainly encodes punishment (the PPL1 cluster) (Saumweber *et al*., 2018; Cognigni *et al.,* 2018; Eichler *et al*., 2017). The functions of the <u>m</u>ushroom <u>b</u>ody <u>o</u>utput <u>n</u>eurons (MBONs) from the various compartments are complex because of extensive interconnections between MBONs and feedback from MBONs back to MBINs. Generally, however, the output from the PPL1-supplied compartments direct avoidance behavior, while that from PAM innervated compartments results in attraction (Thum & Gerber, 2019; Cognigni *et al.,* 2018). The larval mushroom body has ten compartments (Saumweber *et al*., 2018), while the adult has 16 (Aso *et al.,* 2014) (Fig. 1C). In both stages, the PAM cluster neurons innervate primarily the medial lobe compartments, while the PPL1 neurons innervate the vertical lobe and peduncle compartments (Cogneti *et al.,* 2018; Thum & Gerber, 2019).

Output from the compartments is by MBONs that release either acetylcholine, GABA or glutamate. As seen in Fig 1C, in both larva and adult, output from vertical lobe compartments is typically by cholinergic MBONs while that from medial lobe compartments is typically by glutaminergic MBONs. The compartments of the peduncle and base of the lobes have either GABAergic or cholinergic MBONs.

With their shortened embryogenesis, *Drosophila* larvae hatch with only γ neurons. To accommodate the lack of α’β’ and αβ neurons, these neurons grow a larval-specific vertical axon branch to form a vertical lobe. A functional mushroom body, though, also requires the appropriate types of input and output neurons (Tanaka *et al.,* 2008; Aso *et al.,* 2014; Saumweber *et al*., 2018 ). Armstrong *et al*. (1998) used a set of enhancer-trap lines to follow subsets of extrinsic and intrinsic mushroom body neurons through metamorphosis and showed that some neurons functioned in both the larval and adult medial lobes. Our use of a large collection of split-GAL4 lines that express in specific larval MBINs and MBONs (Saumweber *et al.,* 2018) and a conditional flip-switch strategy (Harris *et al.,* 2015) have allowed us to follow the fates of most of the extrinsic cells through metamorphosis. We find that some regions of the mushroom body persist with largely the same components through metamorphosis while other regions are disassembled and rebuilt with new components for the adult.

### Metamorphic fates of the larval MBINs and MBONs

We focused on the uni-compartmental cells with well-defined dendritic or axonal “tufts” that define each larval compartment. Following the convention of Veverytsa & Allan (2013), we make a distinction between neurons that undergo remodeling (but stay in the mushroom body circuit) and those that undergo “trans-differentiation”. We have only morphological criteria for trans-differentiation and have given this designation to any MBIN or MBON that leaves the mushroom body to function elsewhere in the adult brain.

Fig. 2 and Table 1 summarize the fates of the larval MBINs (Saumweber *et al.,* 2018). We lacked suitable lines for the two octopaminergic neurons that innervate the calyx compartment (OAN-a1 &-a2). These cells have a very similar anatomy to the two adult OA-VUM2a neurons (Busch *et al*., 2009), leading us to posit that they are the same neurons. The remaining larval MBINs either remodel, trans-differentiate, or degenerate (Fig 2B,C). Those innervating the contiguous LP, LA, and LVL compartments typically remain as uni-compartmental MBINs and innervate similarly positioned compartments in the adult. The metamorphosis of the octopaminergic MBIN, OAN-g1, is atypical in this group because in its larval form it is a uni-compartmental neuron that innervates only the LVL compartment, while its adult form, known as OA-VPM3, continues to have mushroom body contact in the γ2 compartment but also has extensive arbors in the fan-shaped body and the medial and lateral superior protocerebrum (Busch *et al.,* 2009; Figure 2---figure supplement 1).

**Figure 2.**
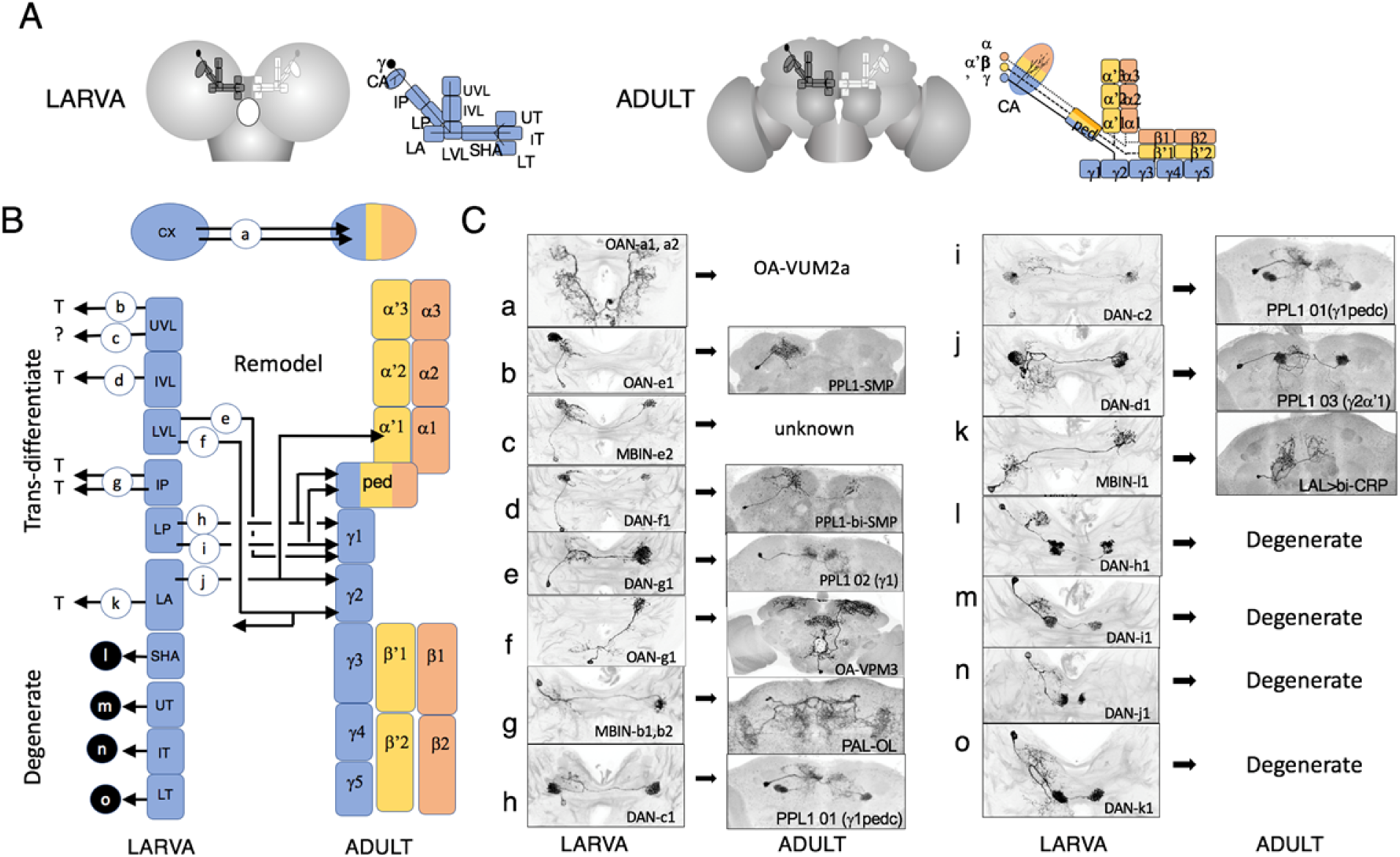
The metamorphic fates of the larval mushroom body input cells (MBINs). **(A)** Comparison of compartment structure of the larval and adult brain. Compartment color indicates the Kenyon cell axons each possesses. Compartment names as in Fig 1. **(B)** Schematic summarizing the fates of the compartmental MBINs of the larva. Arrows indicate whether cells remodel and remain as MBINs, trans-differentiate to have different functions in the adult, or degenerate. Lower case letters refer to images in part C. **(C)** Images comparing the larval and adult forms of the larval MBINs based on flip-switch immortalization. The images and names of the larval cells from Saumweber *et al*. 2018; adult names based on Aso *et al*., 2014 and Li *et al.,* 2020, or this study

**Table 1:**
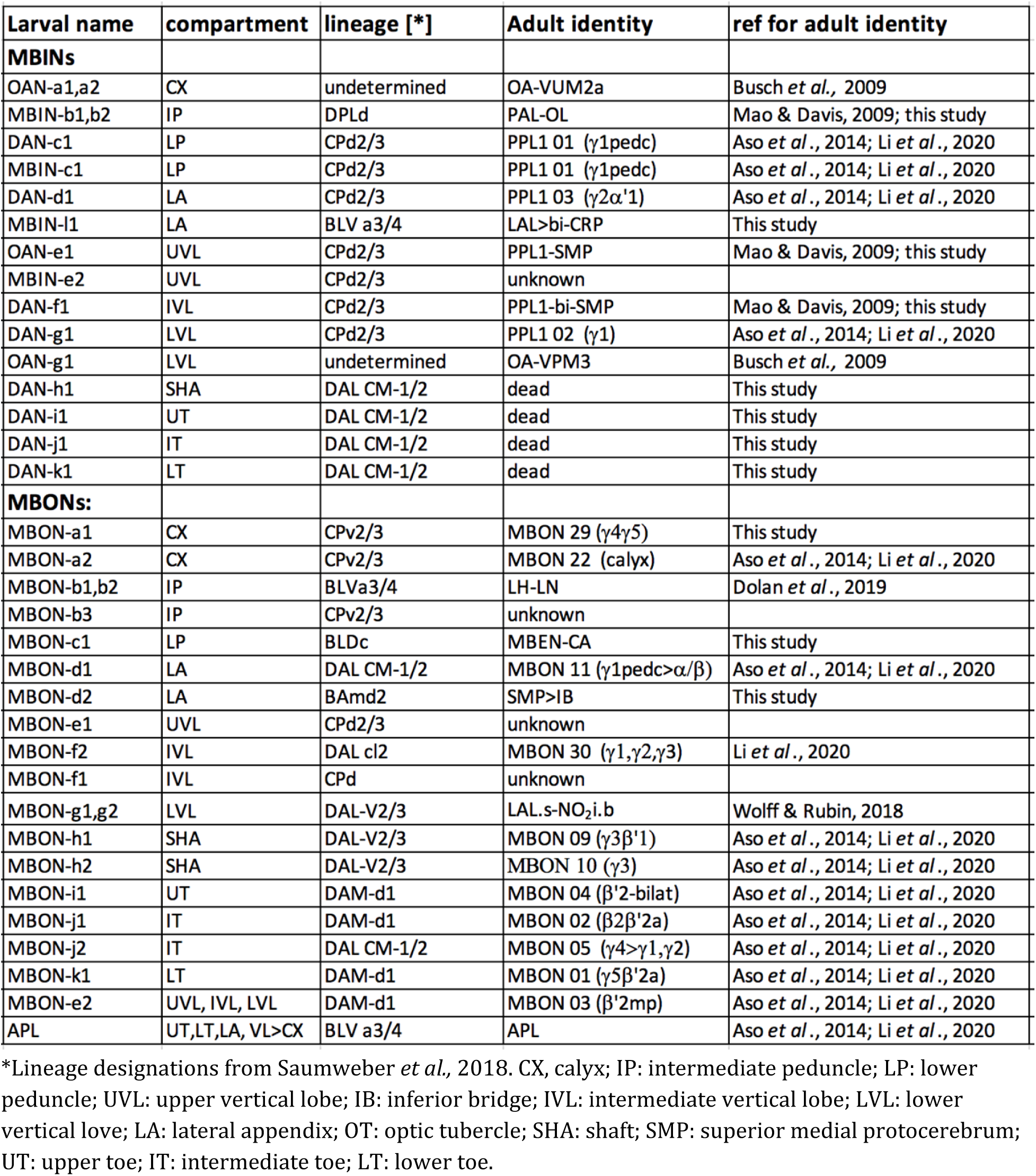
Metamorphic fates of larval mushroom body extrinsic neurons.

The larval MBINs in the more distal compartments of the medial or vertical lobes either trans-differentiated or die. Using our flip-switch approach, we were not able to find adult counterparts for the four PAM cluster neurons innervating the medial lobe compartments (SHA, UT, IT and LT). As shown in the next section, this failure is because they degenerate at the start of metamorphosis. The remaining MBINs, which innervate the larval peduncle and distal vertical lobe compartments trans-differentiate (Figure 2C). The most extreme change was seen for MBINs-b1 and -b2; they withdraw from the upper peduncle of the larva and subsequently become sexually dimorphic neurons that innervate the adult optic lobes (Figure 2Cg; Figure 2---figure supplement 2A,B). In their adult form, they are members of the adult PAL cluster of aminergic neurons (Mao & Davis, 2009) and we have named them PAL-OL. The three vertical lobe MBINs (OAN-e1, MBIN-e2, and DAN-f1) are members of the PPL1 group (Saumweber *et al.,* 2018). We did not have a line that allowed us to determine the fate of MBIN-e2. OAN-e1 reorganizes to innervate the neuropil shell surrounding the mushroom body lobes (as PPL1-SMP; Figure 2---figure supplement 2C) and DAN-f1 forms bilateral arbors in the adult superior medial protocerebrum as PPL1-bi-SMP (Figure 2--- figure supplement 2D). The remaining MBIN is MBIN-l1 which provides input from the lateral accessory lobe to the larval LA compartment, but then trans-differentiates to become neuron LAL>bi-CRP that bilaterally innervates the adult crepine neuropil (Figure 2Bk; Figure 2---figure supplement 2E).

We had lines that allowed us to establish the fates of 14 of the 17 types of larval MBONs (Figure 3; Table 1). None died; the larval cells either remodeled or trans-differentiated. The two cells from the larval calyx (MBON-a1 & -a2) illustrate two extremes of remodeling (Figure 3Ca,b). The larval MBON-a2 neuron assumes a very similar anatomy in the adult where it is called MBON 22 (AKA MBON-calyx). By contrast, the larval MBON-a1 neuron leaves the calyx and is redirected to the γ lobes as the adult neuron MBON 29 (AKA MBON γ4γ5). The larval MBONs innervating peduncle or vertical lobe compartments (IP, LP, UVL, IVL and LVL) either leave the mushroom bodies altogether (Figure 3Cg,h,j) or, like MBON-e2 and -f2, shift to medial adult lobe compartments (Figure 3Cd,e). Our flip-out approach did not give us the adult identity of MBON-c1, which projects from the larval LP compartment, but its early metamorphic outgrowth is directed into the calyx (Fig 3Cj; see next section). By contrast to the MBONs innervating the larval vertical lobe and peduncle compartments, those innervating the medial lobe remain as medial lobe neurons in the adult and supply topologically similar compartments (Figure 3m-r).

**Figure 3.**
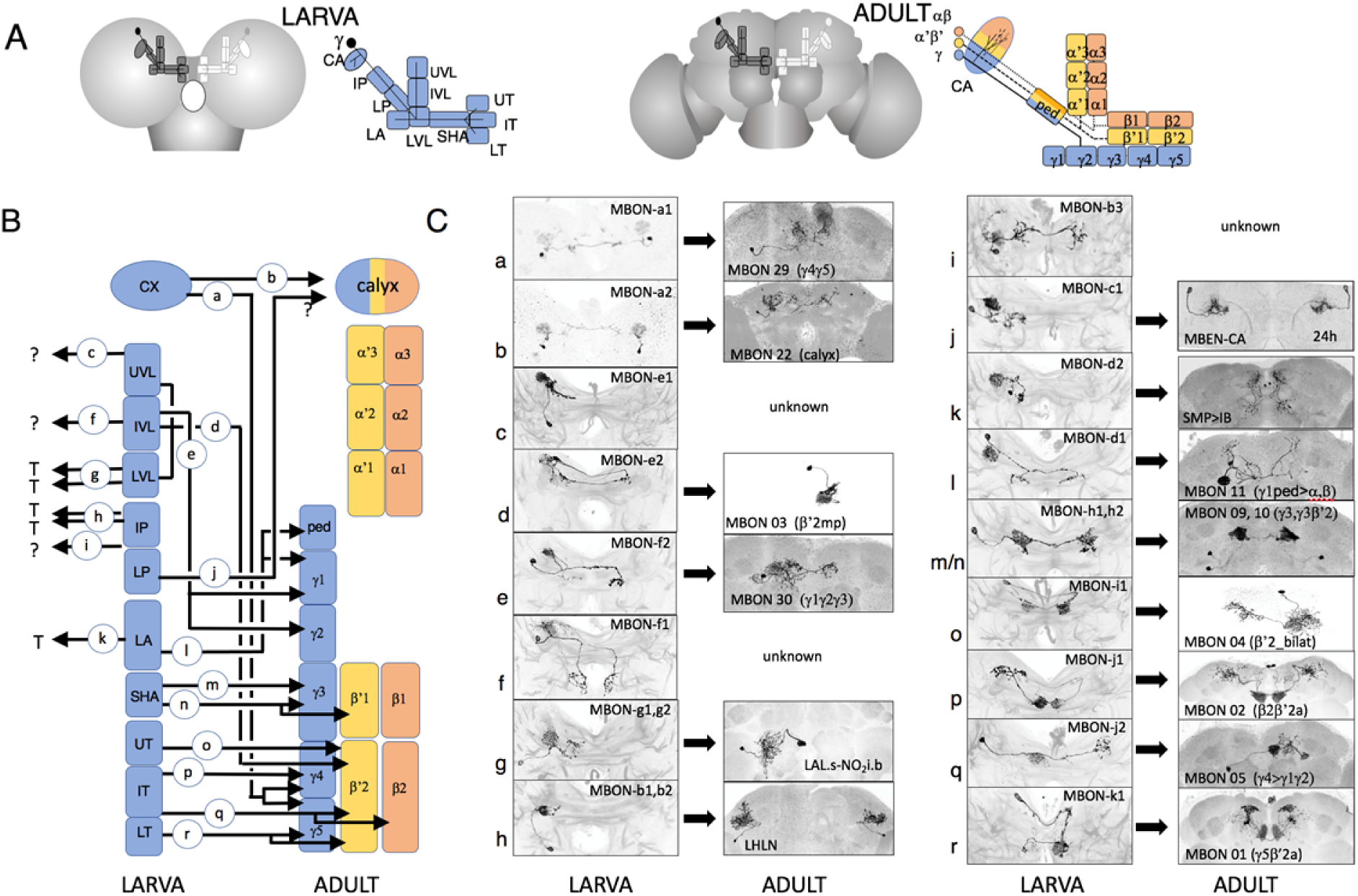
The metamorphic fates of the larval mushroom body output cells (MBONs). **(A)** Schematic summarizing the fates of the compartmental MBONs of the larva. Arrows indicate whether cells trans-differentiate to perform non-mushroom body functions in the adult or remodel as MBONs and project from the indicated adult compartments. Lower case letters refer to images in part C. **(B)** Images comparing the larval and adult forms of the larval MBONs based on flip-switch immortalization. The images and names of the larval cells from Saumweber *et al*. 2018; adult names based on Aso *et al*., 2014 and Li *et al.,* 2020, or this study

The fates of the neurons that undergo trans-differentiation vary markedly. For example, MBON-b1 and -b2 leave the larval IP compartment and become local interneurons in the adult lateral horn (Figure 3Ch; Figure 3---supplementary figure 1B) (LHLN neurons; Dolan *et al*., 2019). The most striking changes occur in MBON-g1 and -g2 (Figure 3Cg; Figure 3---supplementary figure 1C,D), which transform into the LAL.s-NO2i.b neurons that innervate the nodulus of the adult central complex (Wolff & Rubin, 2018).

### The time-course of transitions of larval mushroom body extrinsic neurons

The fates of most of the larval input and output neurons that we determined by the flip-switch method were confirmed by following the expression of the parental lines through early metamorphosis (Figures 4, 5). Many enhancer-based lines change their expression patterns in the transition from larva to adult, but we found that GFP expression typically persists through enough of the remodeling period to confirm the cell’s adult identity.

**Figure 4.**
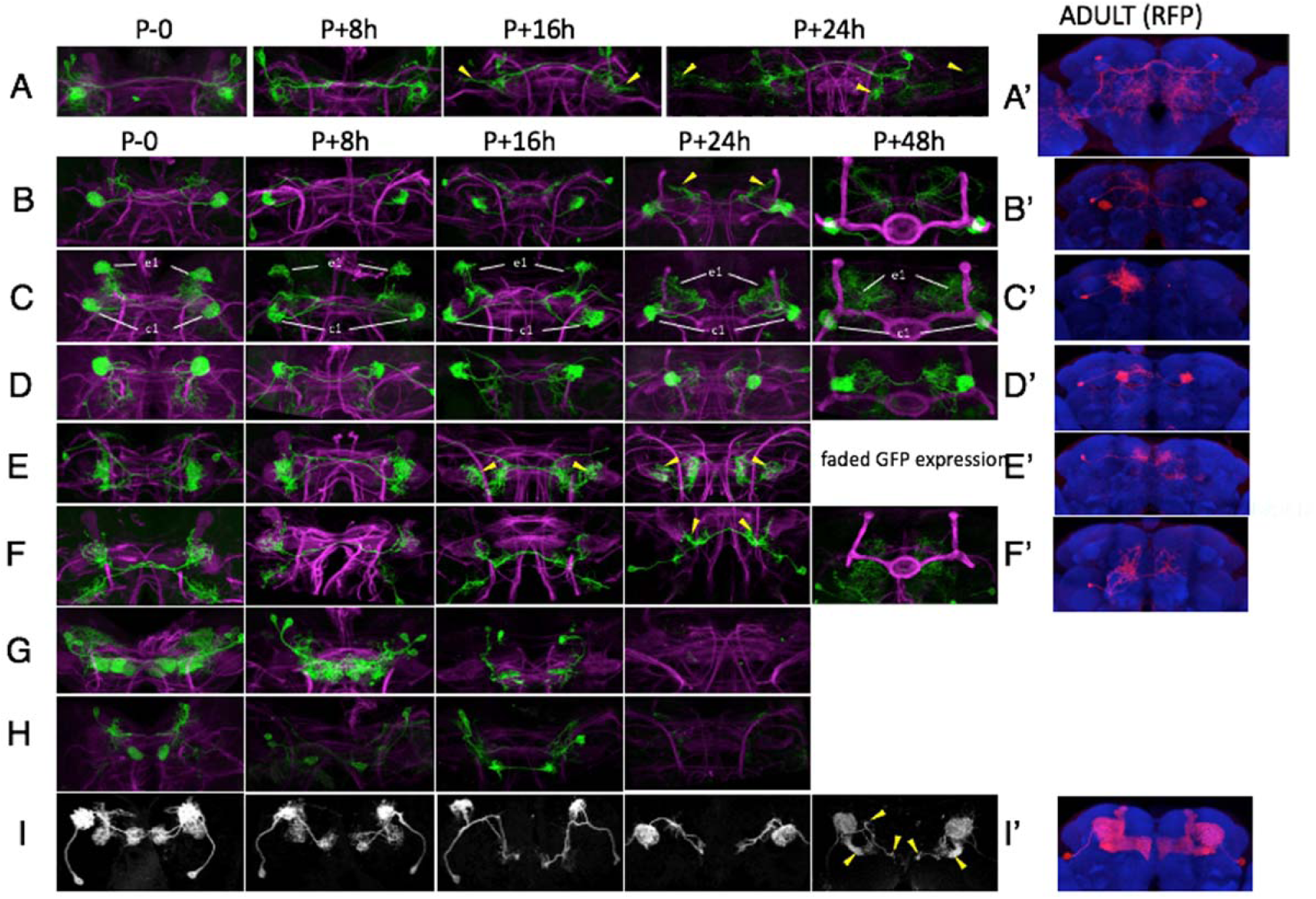
Confocal images of the changes in larval MBINs and APL during the first 48 hr of metamorphosis and their adult appearance (‘). Confocal images show GFP expression through the early hours of metamorphosis; the adult images show flip-switch induced expression of red fluorescent protein (RFP). Arrow heads: growth cones; *: contaminating arbor from other cell types. **(A)** MBIN-b1 & -b2 become PAL-OL interneurons. **(B)** DAN-c1 becomes adult PPL1-γ1pedc . **(C)** OAN-e1 becomes PPL1-SMP; this line also expresses in DAN-c1. **(D)** DAN-d1 becomes PPL1-γ2α’1. **(E)** DAN-g1 becomes PPL1-γ1. **(F)** MBIN-l1 becomes LAL>bi-CRP **(G)** DAN-i1, -j1 and l1 degenerate by 16 to 24 hours after puparium formation (APF). **(H)** DAN-k1 also degenerates. **(I)** APL remodels to become the adult APL. Lines used for developmental timelines: A: JRC-SS21716, B: JRC-SS03066, C: JRC-SS01702, D: JRC-MB328B, E: JRC-SS01716, F: JRC-SS04484, G: JRC-SS-01949, H_ JRC-SS01757, I: JRC-SS01671.

**Figure 5.**
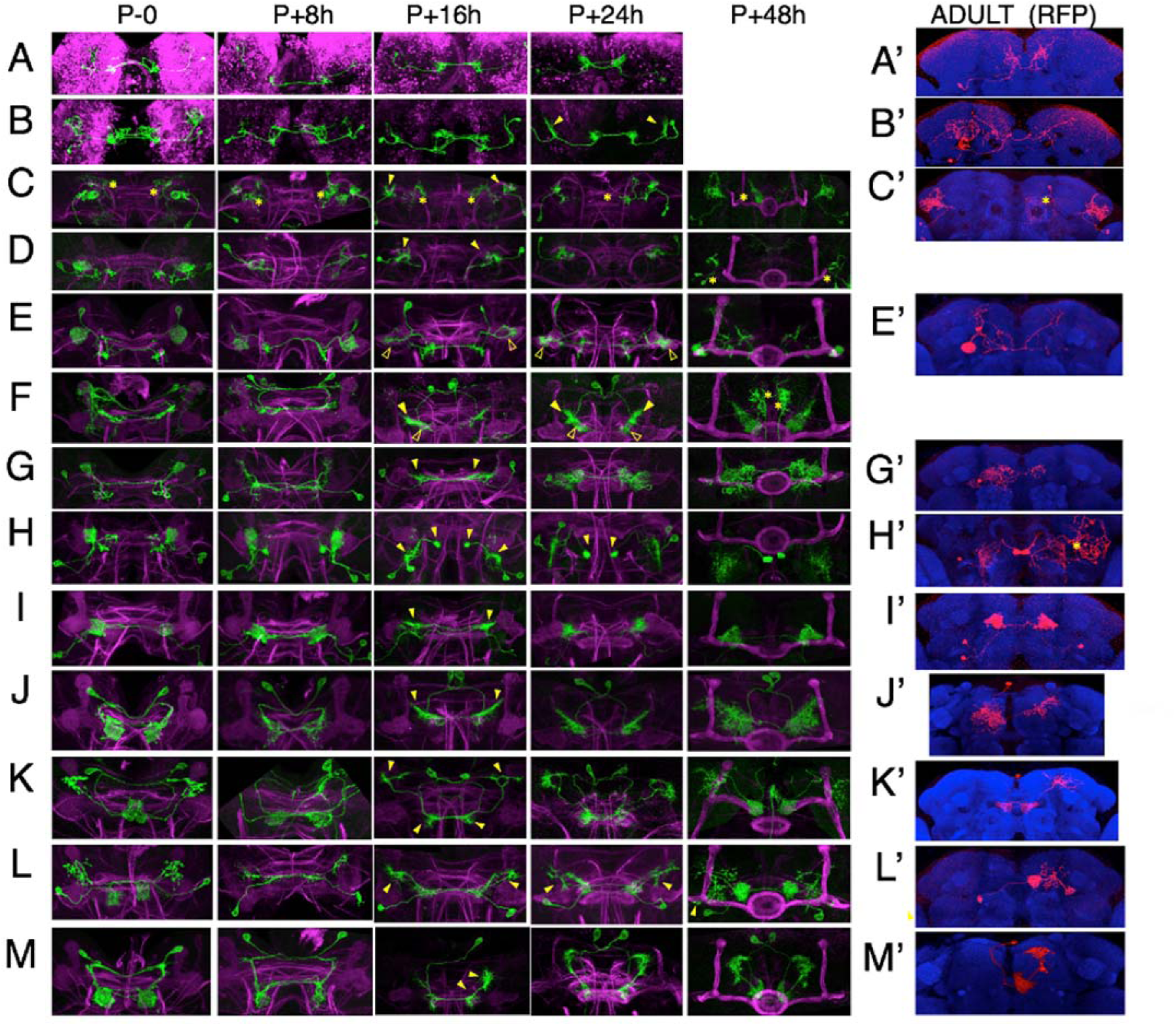
Confocal images of the changes in larval MBONs during the first 48 hours of metamorphosis and subsequent adult appearance (‘). Confocal images show GFP expression through the early hours of metamorphosis; the adult images show flip-switch induced expression of red fluorescent protein (RFP) showing the terminal anatomy of the cells. Arrow heads: growth cones; *: contaminating arbor from other cell types. **(A)** MBON-a1 becomes MBON 29 (γ4, γ5). **(B)** MBON-a1 becomes adult MBON 22 (MBON calyx ). **(C)** MBON-b1 & -b2 become adult lateral horn local neurons. **(D)** MBON-c1becomes MBE-CA. **(E)** MBON-d1 becomes adult MBON 11 ( γ1ped). **(F)** MBON-e2 becomes MBON 03 (β’2mp). **(G)** MBON f2 becomes MBON 30 (γ1γ2γ3). **(H)** MBON-g1 & -g2 become the central complex neurons LAL.s-NO_2_i.b. **(I)** MBON-h1 & -h2 become adult MBON 09 (γ3β’1) and MBON 10 (γ3). **(J)** MBON-i1 becomes MBON 04 (β’2_bilat). **(K)** MBON-j1 becomes MBON 02 (β2β’2a) **(L)** MBON-j2 becomes MBON 05 (γ4>γ1γ2). **(M)** MBON-k1 becomes MBON 01 (γ5β’2a). *Drosophila* lines used for developmental timelines: A: JRC-SS00867, B: JRC-SS02006, C: JRC-SS01708, D: JRC-SS21789, E: JRC-SS01705, F: JRC-SS04559, G: JRC-SS-04320, H_ JRC-SS02130, I: JRC-SS01725. J: JRC-SS04244, K: JRC-SS01973, L: JRC-SS00860; M: JRC-SS01980.

As expected by our failure to find the adult forms of DAN-h1 to -k1 using the flip-switch method, we found that all four neurons degenerated early in metamorphosis (Figure 4G,H). Their dendritic arbors collapsed by 8 hr after puparium formation (APF), cell bodies were disrupted by 18 hr APF, and, by 24 hr APF, they were reduced to scattered GFP-labeled debris.

The remaining MBINs and MBONs either remodeled or trans-differentiated (Figures 4 & 5) and they showed a time-course of pruning and outgrowth that paralleled changes in the γ Kenyon cells. Pruning of γ neuron axons is evident by 4 hr APF and is completed by 16 to 18 hr APF (Watts *et al.,* 2003). Adult outgrowth then commences by 24 hr and is finished by 48 hr (Yaniv *et al.,* 2012; Mayseless *et al.,* 2018). All MBINs (Figure 4) and MBONs (Figure 5) showed arbor loss by 8 hr APF and were completely pruned by 16 hr APF. Growth cones became evident between 16 and 24 hr APF. MBIN-b1 & -b2 showed the most exuberant outgrowth, having formed growth cones that were halfway to the optic lobes by 16 hr APF and reaching these structures by eight hours later (Figure 4A and supplementary figure 1). Most neurons achieved their adult form by 48 hr APF. The APL and MBON-j2 neurons were exceptional in their outgrowth because they continued arbor extension beyond 48 hr (Fig. 4I and 5L). APL eventually covers all of the adult compartments by 72hr APF (Mayseless *et al.,* 2018) and provides inhibitory feedback from the lobes to the calyx. For MBON-j2, its γ4 tuft formed at the same time as those of other MBONs but the formation of its γ2 and especially its γ1 arbors was delayed. (Figure 5---supplement figure 1). In its adult function as MBON 05 (AKA MBON-γ4>γ1,γ2), it provides feed-forward inhibition from γ4 to γ2 and γ1 (Aso *et al.,* 2016). Its extended period of outgrowth may allow time for the compartment microcircuits to become established before it interconnects them.

MBON-a1 and -a2 provide an interesting contrast of divergent remodeling of two similar larval cells (Figure 5A,B). Both cells were retracting their dendritic and axonal arbors by 8 hr APF. MBON-a1 completely removed its dendritic arbor by 16 to 24 hr, and the distal portion of the neuron extended new growth to innervate the γ4 and γ5 compartments and surrounding neuropils. In contrast, the dendritic arbor of MBON-a2 only partially regressed, organized into a dendritic growth cone by 16 to 24 hr APF, and reinvaded the calyx. In its adult form as MBON-20 (MBON-calyx; Aso *et al.,* 2014; Li *et al.,* 2020), its dendritic and axonal arbors are very similar to its former larval form.

The most stable extrinsic neuron is DAN-d1. Its dendritic arbor remained intact through metamorphosis and its axonal tuft underwent a mild, transient thinning from 8 to 18 hr APF and then extended fine processes into the forming γ2 and α’1 compartments between 24 and 48 hr APF (Figure 4D). A more extreme remodeling was evident in MBON-d1 (Figure 4E). Its larval dendritic tuft retracted and reorganized into a growth-cone by about 18 hr APF, extended into the γ1 and peduncle compartments by 24 hr, and achieved its adult dendritic configuration by 48h. Its axonal arbor expanded from a compact projection in the larva to a sparse arbor along the adult α and β lobes (Aso et al., 2017; Li et al., 2020).

Amongst the cells undergoing trans-differentiation, the most extreme changes were seen in MBON-g1 & g2, which shifted from the larval mushroom body to the adult central complex (Figure 5H). Their larval axonal arbors were retracting by 8 hr APF and organized into an axonal growth cone by 18 hr APF. The growth cone then navigated medially to innervate the intermediate zone of the nodulus. The dendritic tufts of MBON-g1 & -g2 thinned by 8 hr APF and then fragmented as new dendritic growth cones sprouted from the base of the old arbor (18 hr APF). The dendrites invaded and expanded to cover the ipsilateral lateral accessory lobe. A similar level of extreme change between larval and adult morphologies was seen for MBIN-b1 and b2 that became sexually dimorphic, optic lobe input interneurons in the adult (Figure 2 – figure supplement 2A,B).

MBON-c1 was refractory to the flip-switch approach, but its early metamorphic changes gave us insight into its adult function. As seen in Figure 5 – figure supplement 2, by 24 hr APF its larval dendritic arbor was essentially gone, and new growth cones have invaded the calyx. This line lost its GFP expression after this time but the extensive growth into the calyx by 24 hr indicates that this neuropil is its terminal, adult target. Its anatomy at 24 hr APF, though, was too immature to match it to any known adult cell, so we have called its adult form MBEN-CA (mushroom body extrinsic neuron to calyx).

### The larval form is a derived state for the neurons that show trans-differentiation

For trans-differentiating neurons, that assume two different identities, is one identity ancestral for the cell and the other identity derived to accommodate metamorphosis? Establishing such homologies is difficult for the extrinsic neurons but it is possible for neuron groups in the VNC, such as the midline spiking interneurons (Figure 6). These neurons are found in a large cluster in each thoracic hemineuropil and each cluster is the progeny of a single, identified neuroblast (Shepherd & Laurant, 1984). Their cell bodies lie near the ventral midline and they project to the contralateral leg neuropil where they shape the response of leg motoneurons to input from leg mechanoreceptors (Siegler & Burrows, 1984). They were first described in grasshoppers (Siegler & Burrows, 1984) but they are found in both direct developing and metamorphic insects (Witten & Truman, 1998), indicating an involvement in leg circuitry through insect evolution.

**Figure 6.**
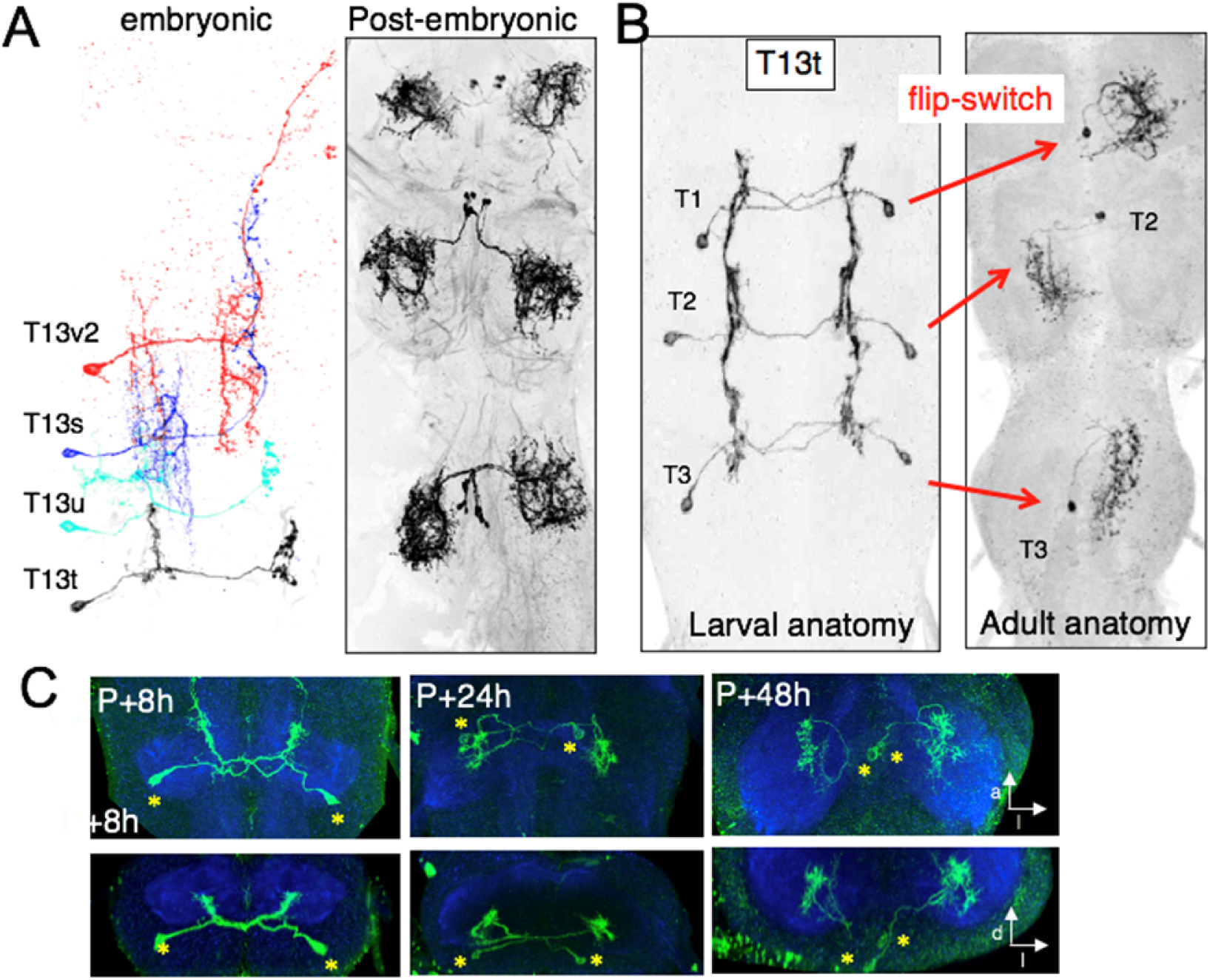
Form and fates of lineage 13 (NB4-2) thoracic interneurons. **(A)** (Left) staggered images of four different lineage 13B larval interneurons in the segment T3; (right) example of the form of the postembryonic-born neurons from the same lineage. (**B)** (Left) image of the larval form of T13t neurons revealed by the JRC-SS04274 driver line and (right) examples of the terminal form of these neurons in the adult as revealed by the flip-switch method. (**C)** Dorsal (top) and transverse (bottom) views of the early metamorphosis of the T3 pair of T13t cells: the dendritic arbor is gone by 8 hours after pupariation (P+8h) , contralateral growth cones are evident by P+24h, and the arbor is near its maximal extent by P+48h. Through this period, the expanding neuropil pulls the cell bodies (*) to their adult position near the midline.

In *Drosophila*, these inhibitory neurons arise from the neuroblast NB4-2 during the postembryonic phase of neurogenesis (Harris *et al.,* 2015; Lacin & Truman, 2016). They are stockpiled as immature cells during larva growth and then mature into midline inhibitory interneurons at metamorphosis. The embryonic-born neurons of this lineage (e.g. neurons T13s, T13t, T13u and T13v2; Figure 6A), however, are commissural interneurons that look nothing like their postembryonic siblings. However, as shown in Figures 6B and C, at metamorphosis these neurons trans-differentiate to assume the same terminal identity as the postembryonic-born cells. Therefore, we see that just like in the direct developing grasshopper (Shepherd & Laurant, 1984), *Drosophila* starts producing its midline leg interneurons during embryogenesis but the early-born cells initially take on an **assumed** identity adapted to larval needs and defer expressing their ancestral identity as leg interneurons until metamorphosis. This comparison between direct developing and metamorphosing insects, then, argues that the adult identity of a trans-differentiating neuron represents its ancestral identity while its assumed identity in the larva is a derived identity that evolved to accommodate the highly modified larval stage. The significance of this relationship will be considered below.

### Origins of adult-specific Mushroom Body input and output neurons

Remodeled larval MBINs and MBONs do not account for all of the input and output neurons of the adult mushroom bodies (Figures 2B and 3B). The adult has about 40 different types of MBINs and MBONs (Aso *et al.,* 2014; Li *et al.,* 2020) and we found that only 15 types also function in the larval structure. The remaining 25 types could either come from other trans-differentiating neurons whose larval function is outside of the mushroom bodies or they could be born during the postembryonic neurogenic period. The origins of 22 of these 25 types are summarized in Figure 7A,B and Table 2. They are all postembryonic born. We found no neuron whose terminal fate was as a mushroom body extrinsic neuron but assumed a different function in the larva.

**Figure 7.**
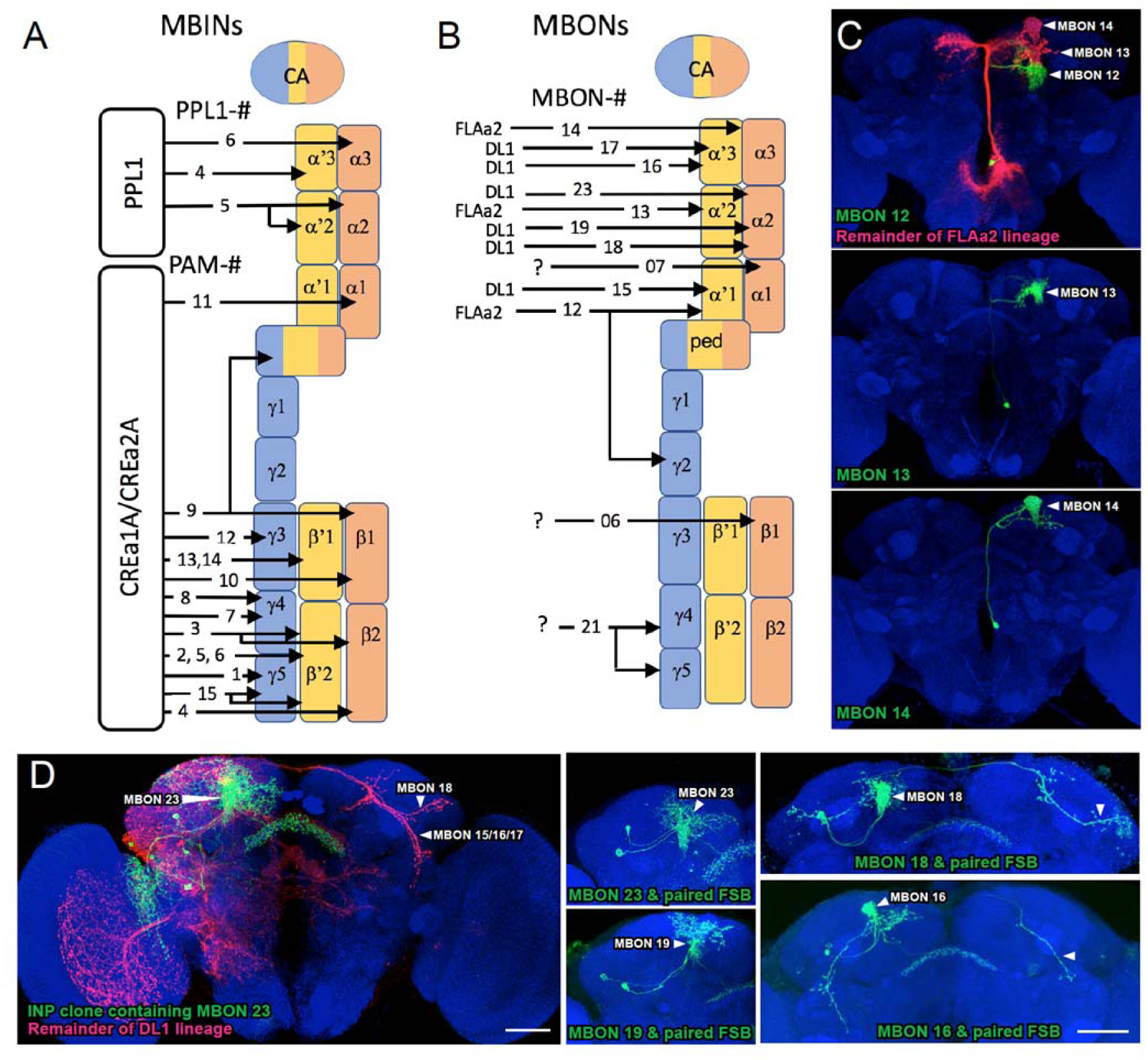
The Postembryonic-born MBINs and MBONs of the adult mushroom bodies. **(A)** Summary of the origins of to the various compartments for the adult mushroom bodies. Numbers give the number of neurons in each of the PAM groups (from Aso *et al.,* 2014). **(B)** Summary of the postembryonic-born MBONs added to the adult mushroom bodies from the indicated lineages. **(C)** Results of twin-spot MARCM approach showing the sequential postembryonic birth of different neurons from the FLAa2 lineage which innervate of some the α’ and α compartments. C, C’ and C” images are produced by successively later heat-shocks in the larva; green cells are produced after the heat-shock while the red cells (shown only in C) are the remainder of the lineage. **(D).** Twin spot MARCM results from type II DL1 lineage. The leftmost panel shows the progeny of an intermediate neural progenitor (INP) in green and the remainder of the lineage in green. The remaining panels show GMC clones with an MBON neuron and its paired sister fan-shaped body (FSB) neuron.

**Table 2:**
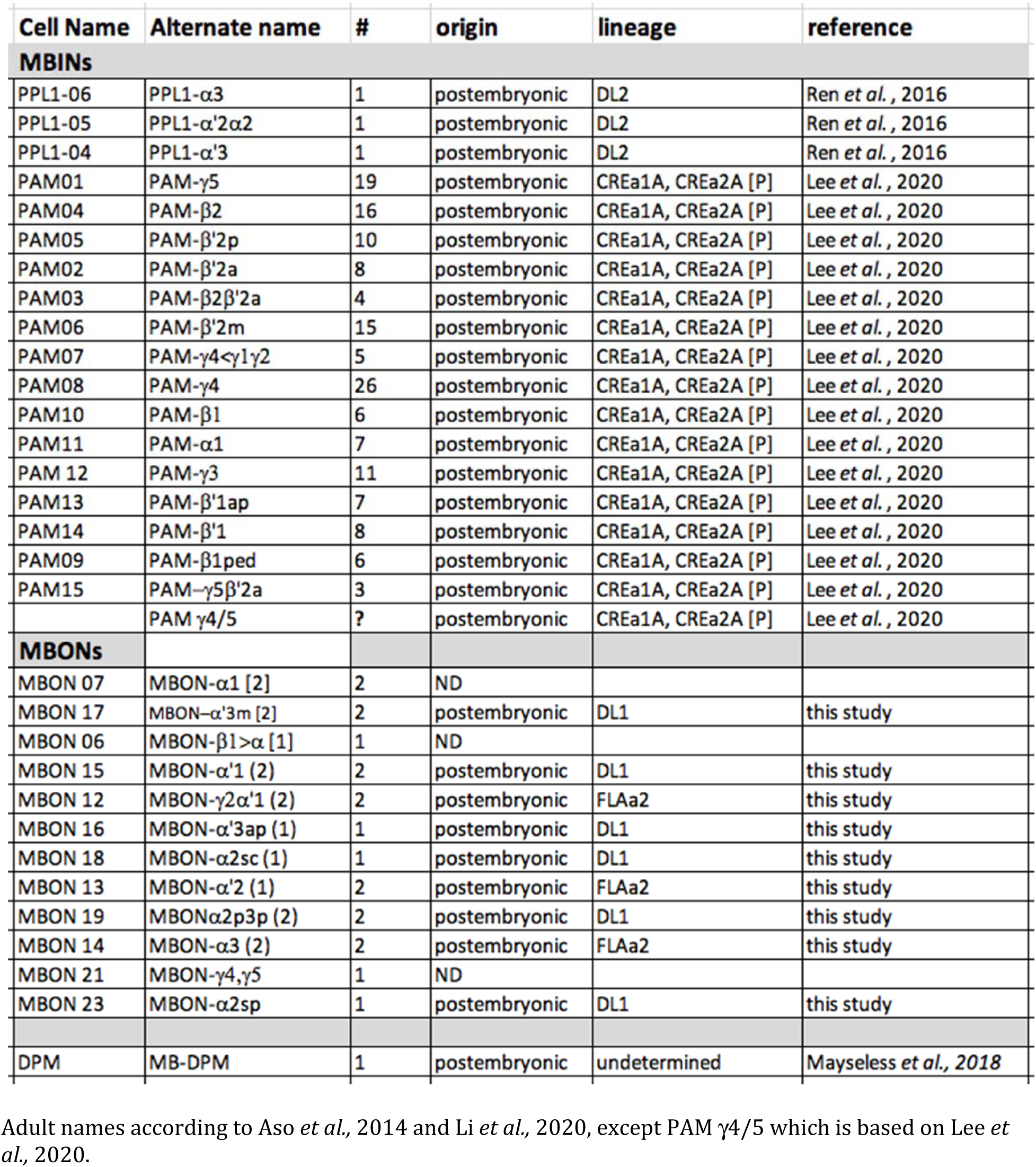
Developmental origins of adult MBINs and MBONs that do not come from remodeled larval, extrinsic mushroom body neurons.

**Table 3 :**
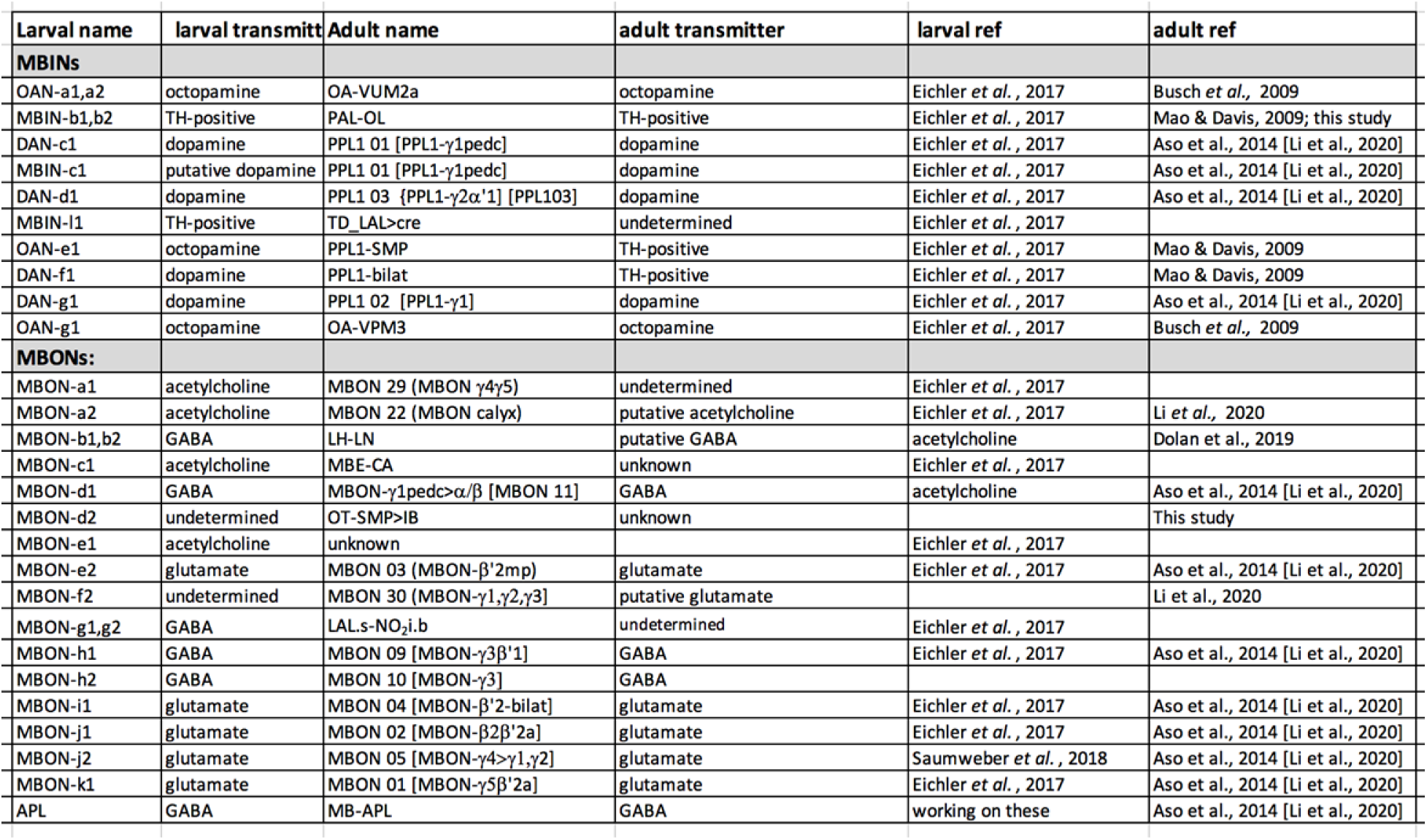
Comparison of transmitter expression in larval and adult forms of MBONs and MBINs.

The MBIN side of the adult circuit is dominated by the addition of about 150 PAM-class neurons divided into at least 15 different types (Aso *et al.,* 2014; see also Lee *et al.,* 2020; Li *et al.,* 2020). These neurons are born during the postembryonic phase of neurogenesis and come from the CREa1A and CREa2A lineages (Lee et al., 2020) (Table 2). This massive addition of PAM cluster neurons suggests an enhanced sophistication for discriminating rewarding stimuli in the adult versus the larva (*e.g., Li* et al., 2020). By contrast, there is no change in the number of PPL1 neurons that primarily provide input to the compartments involved in aversive conditioning. While the number remains constant, though, the cell population partially changes: the adult adds three new PPL1 neurons to the set (Ren *et al.,* 2016), but loses three larval PPL1 neurons to other brain circuits of the adult (Figure 2).

For the MBONs, the origins of nine of the twelve remaining types (Table 2; Fig 7B) were determined using the twin-spot MARCM technique (Yu et al., 2009). As seen in Fig. 7C, MBONs 12 (also called MBON-γ2α’1), 13 (MBON-α’2), and 14 (MBON-α3) are born in succession during the postembryonic divisions in the FLAa2 lineage. Most of the remaining vertical lobe MBONs arise in the DL1 lineage (Fig 7D; Table 2). DL1 is a Type II neuroblast that has an atypical division pattern (Boone & Doe, 2008; Wang *et al*., 2014). Each neuroblast division produces an intermediate precursor cell which, in turn, produces a small series of ganglion mother cells. Each of the latter divides to make two daughter neurons. As shown in Figure 7D for a number of these ganglion mother cells, one daughter becomes a vertical lobe MBON while the other becomes a central complex neuron, innervating the fan-shaped body.

## Discussion

### Strategies for generating neurons for the larval CNS

Our analysis of the metamorphosis of the mushroom body has revealed at least five strategies for producing larval neurons (Figure 8A). The **first** strategy is that a neuron acquires its terminal identity during embryogenesis. It has similar form and function in both larva and adult although there may be some remodeling of arbors going from one stage to the other. DAN-d1, for example, shows little change from larva to adult. Its dendritic arbor remains largely intact through metamorphosis while its axonal tuft shows only a moderate thinning before re-expanding (Figure 4D). MBON-j1 provides a more typical pattern. Its larval and adult forms are quite similar, but the larval cell still goes through severe axonal and dendritic pruning but then regrows into a very similar morphology for the adult (Figure 5K).

**Figure 8.**
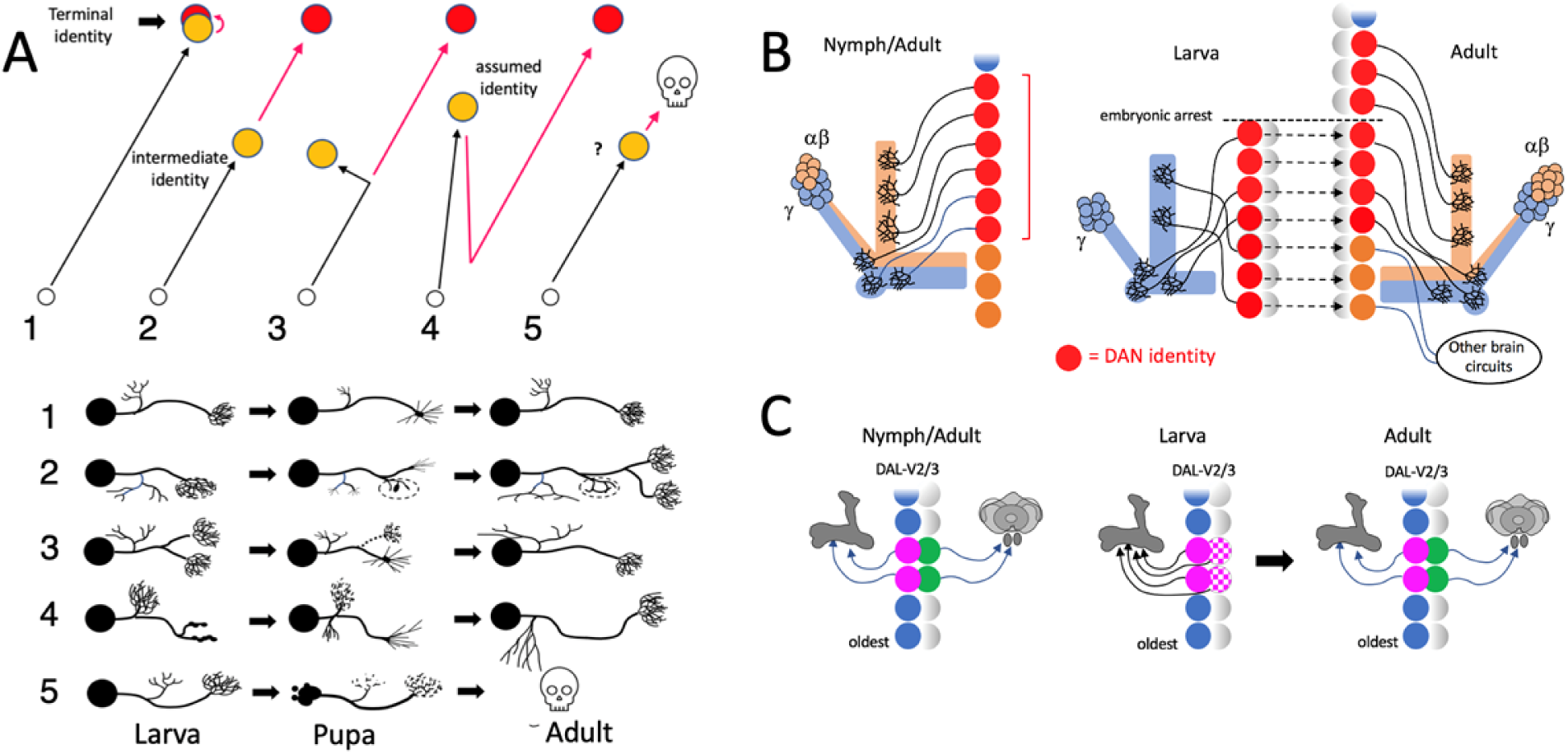
Cellular strategies for producing larval-adapted neurons. **(A)** Schematic comparison of larval (yellow) and adult (red) forms of neurons to illustrate strategies for producing larval cells. The red portion of the path occurs during metamorphosis: (1) neuron acquires its terminal identity by hatching and serves similar roles in larva and adult , (2) larval form is based on an intermediate phase in the development of the neuron’s terminal identity. (3) larval and adult cells are similar but larval cell develops significant features that are lost in the adult, (4) the neuron undergoes trans-differentiation in which the larval form has a assumed identity that differs from its adult identity, (5) recruitment of neurons for transient survival in the larva from the populations of neurons that normally die during embryogenesis. **(B,C)** Potential mechanisms for allowing a trans-differentiating neuron to assume a temporary identity. **(B)** Generating trans-differentiated PPL1 DAN neurons in the DL2 lineage involves modifying the temporal program that establishes neuron identities within its lineage. In direct developing insects, the PPL1-DANs arise during embryogenesis, as a temporal block of neurons consisting of one daughter cell from each of a sequential series of neuronal precursor cells (red neurons). In *Drosophila,* only some of these neurons are produced before the embryonic neurogenic arrest, but more neurons with the appropriate phenotype are made by temporarily expressing the appropriate determinants in other early-born cells in the lineage. At metamorphosis, these determinants are lost and replaced by ones (orange) characteristic for their terminal identity. **(C)** Hypothetical scheme in which identity differences between sibs is used to produce temporary larval MBONs in the DAL-V2/3 lineage. In direct developing insects two sequential neuronal precursors are proposed to divide to produce one daughter directed to the mushroom bodies and the other to the central complex. In larvae of *Drosophila,* the second daughter cell temporarily assumes some of the features of its sibling and re-targets to the larval mushroom body. It assumes its proper central complex role at metamorphosis.

The pruning of larval neurons at metamorphosis is caused by a global, hormonal signal -- the ecdysone surge that drives metamorphosis (Lee *et al.,* 2000; Schubiger *et al*., 1998). Local, extrinsic factors, though, can also influence the extent of pruning (Williams and Truman, 2005). The importance of such factors for pruning in the mushroom body circuit was shown by studies on the APL neuron (Mayseless *et al.,* 2018). APL pruning is dramatically suppressed by the selective inhibition of axon degeneration in the γ Kenyon cells, the major synaptic partner of this neuron. The remodeling of MBINs may have a similar dependence on the state of their postsynaptic targets. The γ Kenyon cells prune back their axons from the distal compartments but retain some processes in the LP and LA compartments at the base of the lobes. In parallel, the MBINs to the latter two compartments (i.e., DAN-c1 and -d1) show only moderate pruning (Figure 4B,D: P+16h and P+24h), while those innervating more distal compartments prune severely and regrow their arbors from defined growth cones (*e.g.,* Figures 4E,F). This pattern suggests that the severity of axon pruning by the MBINs depends on the extent of loss of their postsynaptic targets.

The quantitative differences seen in the MBINs, though, are not seen in the MBONs. The latter undergo severe dendritic pruning and regrow from growth cones, regardless of which compartment they innervate (Figure 5). This difference between MBONs and MBINs suggests that dendrite pruning may be less dependent on the state of local environments as compared to axonal pruning.

The **second** strategy is that a neuron’s larval form is based on the cell arresting its development at some intermediate point in its trajectory to its terminal identity. This is analogous to the case of ingrowing thalamic neurons in the developing mammalian forebrain (Kanold *et al*., 2003). These early-born neurons arrive prior to the birth of the cortical granule cells so they initially synapse on subplate neurons. After the granule cells are born, they lose their connections in the subplate and move into the cortex. A similar scenario in *Drosophila* is suggested by the two sets of octopaminergic cells, OAN-g1 and OAN-a1 & -a2. We find that the adult form of OAN-g1 is OA-VPM3 (Fig 2Cf; Busch *et al.,* 2009; Busch & Tanimoto, 2010).

Its adult form has a minor contact with the γ2 compartment and then branches exuberantly in the fan-shaped body of the central complex and in the superior medial protocerebrum. The larval form of the cell, though, stops in the LVL compartment (the larval homolog of γ2) and only continues on to the central complex and superior medial protocerebrum at metamorphosis. A similar situation is seen for OAN-a1 & -a2. In their larval form they innervate the antennal lobes and mushroom body calyx (Figure 2Ca), but their terminal form in the adult extends beyond the mushroom body to the lateral horn (Busch *et al.,* 2009).

The **third** strategy is illustrated by larval neurons that acquire a functionality that is absent from their terminal identity. The γ Kenyon cells provide an excellent example in that their larval form possesses a vertical axon branch that is lacking in their adult form. Importantly, this vertical branch is also absent from γ neurons in species that do not have a larval stage (e.g., the cricket; Malaterre, *et al.,* 2002) supporting the idea that it is a larval-specific modification.

The **fourth** strategy is trans-differentiation, which provides the neuron with two distinct identities: its **terminal identity,** which is evident in the adult, and an **assumed identity,** which is seen in the larva. The most extreme example which we found involved the two pairs of central complex neurons (LAL.s-NO_2_i.b) that assumed the identities of mushroom body neurons in the larva (as MBON-g1 and -g2; Fig. 5H). We also found adult optic lobe neurons (Figure 4A) and lateral horn neurons (Figure 5C) that first assumed identities as a mushroom body extrinsic neurons in the larva. A less extreme identity shift was seen for PPL1-SMP. In its larval form as OAN-e1, it innervates the Kenyon cell core of the UVL compartment but in its terminal identity it shifts to the neuropil shell surrounding the vertical and medial lobes (Figure 4C). Trans-differentiation of neurons through ontogeny is not unique to insect metamorphosis. It is also seen in developing zebrafish in the case of dorsal root ganglion sensory neurons that transform into sympathetic neurons after they migrate away from the dorsal root ganglion (Wright *et al*., 2010).

We do not know if there is a qualitative distinction between cells that remodel versus those that trans-differentiate or if these just represent the two extremes of a continuum. Some cells, like MBON-e2 and MBON-j2, have features of both strategies. They function as MBONs in both larva and adult but their roles within the mushroom body circuitry change. MBON-j2 provides output from a medial lobe compartment to contralateral brain regions outside of the mushroom bodies while its terminal identity is to provide intercompartmental communication with the γ lobe (Figure 5L). MBON-e2 makes the opposite shift. Its assumed function in the larva is to assemble output from multiple vertical lobe compartments while its terminal function in the adult is as a uni-compartmental MBON in the adult β’2 compartment (Figure 5F).

The **fifth** strategy is for larval neurons to be recruited from the population of neurons that normally die during embryogenesis. This is a common strategy in the VNC. Neurons are generated according to a reiterated segmental plan in the VNC, but in direct developing insects like grasshoppers, many neurons needed in the thorax are not needed in the abdomen. They are nevertheless produced in both regions, but segment-specific cell death then removes the extraneous cells from the abdomen (*e.g.,* Thompson & Siegler, 1993). Larvae, like those of *Drosophila*, that use abdominal-based crawling for locomotion, though, require many more abdominal neurons than survive in the abdominal CNS of a grasshopper nymph. These larvae appear to acquire needed neurons by drawing from this large pool of embryonic abdominal neurons that are fated to die (Truman, 2005). Their death is delayed so that they can function in the larva but once the larval stages are finished, they revert to their ancestral fate and degenerate. This use of such phylogenetically “doomed” neurons is common in the abdominal segments but less prevalent in the larval thorax or the brain. Indeed, of the larval brain neurons in this study, only the four PAM neurons die after the larval stage is completed (Figure 4G,H). It is curious that these four neurons die rather than remodeling to join the ∼150 new PAM neurons that are added for the adult (e.g., Lee *et al*., 2020). It may be that “doomed” neurons such as these are able to assume a temporary identity that allows them to survive and function in the larva but must then revert to their terminal fate of degeneration at metamorphosis.

### Factors related to trans-differentiation

How are neurons selected to undergo trans-differentiation and assume identites for use in the larval mushroom body circuitry? Besides the observation that many of these neurons are fated for roles in adult-specific circuits, we find that five of the seven types of trans-differentiating neurons come from lineages that produce other neurons that are terminally fated to function in the mushroom bodies (Table 1). The OAN-e1 and DAN-f1 neurons that function in the mushroom body only in the larva come from the CPd2/3 lineage that also produces the PPL1 01 (= larval DAN-c1), PPL1 02 (= larval DAN-g1) and PPL1 03 (= larval DAN-d1) neurons that function in both larval and adult mushroom bodies. The larval MBIN-l1 and MBON-b1, -b2 neurons come from the BLVa3/4 lineage that produces APL, and the larval MBON-g1 and -g2 neurons come from the DAL-V2/3 lineage that produces MBON 09 and 10 ( = larval MBON-h1 and -h2) (Table 1; Saumweber *et al*., 2018).

Two main mechanisms are involved in establishing neuronal identities within a lineage: are established by: one is the relative timing of birth of the neuron’s ganglion mother cell (Kohwi & Doe, 2013; Doe, 2017; Miyares & Lee, 2019) and the other is Notch-dependent signaling between the two siblings from the ganglion mother cell division (Skeath & Doe, 1998). As summarized in Fig. 8B,C, we speculate that either of these mechanisms could be exploited to recruit neurons for temporary function in the mushroom bodies.

The trans-differentiation of a subset of the PPL1 neurons (Figure 8B) could involve modifying temporal fate determination within the PPL1 cluster. The larva has seven tyrosine hydroxylase positive neurons in its PPL1 cluster (Selcho et al., 2009) and these innervate the core of the larval mushroom body. The adult has twelve neurons in this cluster but, again, only seven of these innervate its mushroom body core while the remainder target surrounding neuropils (Mao & Davis, 2009). We find, though, that the seven neurons that innervate the mushroom body are not all the same in the two stages. The three neurons supplying the larval vertical lobe leave the mushroom body circuit and join other brain circuits of the adult. They are replaced by three new PPL1 neurons that are added during the second neurogenic period in the larva (Ren *et al.,* 2016). Two neuroblasts contribute to the PPL1 cluster in both the larva (Saumweber *et al,* 2019) and the adult (Ren *et al.,* 2016). We assume that those innervating the adult mushroom body likely come from the same neuroblast, although only four of them are born during embryogenesis and can therefore contribute to the larval circuit. The missing three neurons either come from the other stem cell or by the temporary transformation of other early-born cells in the DL2 lineage (Figure 8B). Although such genes have not been found for these neurons, we assume that there are terminal selector genes (Hobert & Kratsios, 2019) that determine the phenotype of the PPL1 neurons that innervate the mushroom body. We speculate that the appropriate selector genes may be expressed prematurely in the DL2 lineage to allow other cells to assume an MBIN identity. For these cells, though, the MBIN selector gene expression is temporary and lost at metamorphosis, thereby allowing the neurons to switch to their ancestral fate in the adult.

In contrast to the above scenario, the ability of the central complex neurons, LAL.s-NO_2_i.b, to temporarily assume identities as the larval MBON-g1 and -g2 may involve suppressing the divergent fates of sibling neurons as depicted in Figure 8C. The two neurons arising from the division of a ganglion mother cell typically have very distinct terminal identities with *Notch* signaling determining the difference between the “A” (Notch-on) and the “B” (Notch-off) phenotypes (Skeath & Doe, 1998). *Hey* (Hairy/enhancer-of-split like with a Y) is a bHLH-O protein which is an important Notch target for establishing the “A” phenotype (Monastirioti *et al.,* 2010). Interestingly, *Hey* expression is not *Notch* dependent in the Kenyon cell lineage (Monastirioti *et al.,* 2010), and in these lineages the two sibling neurons are identical. We find that sibs that show central complex versus mushroom body fates show up multiple times in the DL1 lineage (Figure 7D). We speculate that the MBON 09/10 versus LAL.s-NO_2_i.b phenotypes seen in the DAL-v2/3 lineage represents a similar divergence of sibling phenotype with one destined for the central complex and the other for the mushroom body. Alteration of Notch signaling in the embryo, though, might allow the neuron destined for the central complex to temporarily assume characteristics of its mushroom body sibling, thereby becoming another MBON. At metamorphosis, the latter cell would somehow reverse the altered Notch effects and acquire its terminal identity as a central complex neuron. The scenarios in the last two paragraphs are quite speculative but suggest possible avenues of research that could be explored to determine the basis for trans-differentiation.

In some of the cases we examined, the interneurons that acquired an assumed identity in the larva lacked the targets that are appropriate for their terminal identities. In contrast to the interneurons, the embryonic born motoneurons that lack their adult targets do not seem to have the plasticity to assume other functions. The indirect flight motoneuron, M5, (Consoulas *et al*. 2002) and the embryonic-born leg motoneurons of lineage 15 (Lacin & Truman, 2016) lack their normal adult targets in the larva and they remain in an arrested immature state through larval growth and delay their maturation until metamorphosis. A similar developmental arrest has yet to be found amongst embryonic-born interneurons. Given the small number of interneurons that are available to it, the embryo likely uses every interneuron that is available to make its larval CNS.

### Relationship of cell fate to mushroom body compartments

Based on the analysis of enhancer trap lines, Armstrong *et al.,* (1998) found that larval extrinsic neurons mainly ended up in the γ lobe compartments of the adult. As seen in Figure 9A, our cell-by-cell analysis of the fates of the larval neurons reached a similar conclusion with the additional insight that some of the larval cells leave the mushroom bodies at metamorphosis and function elsewhere in the adult brain.

**Figure 9:**
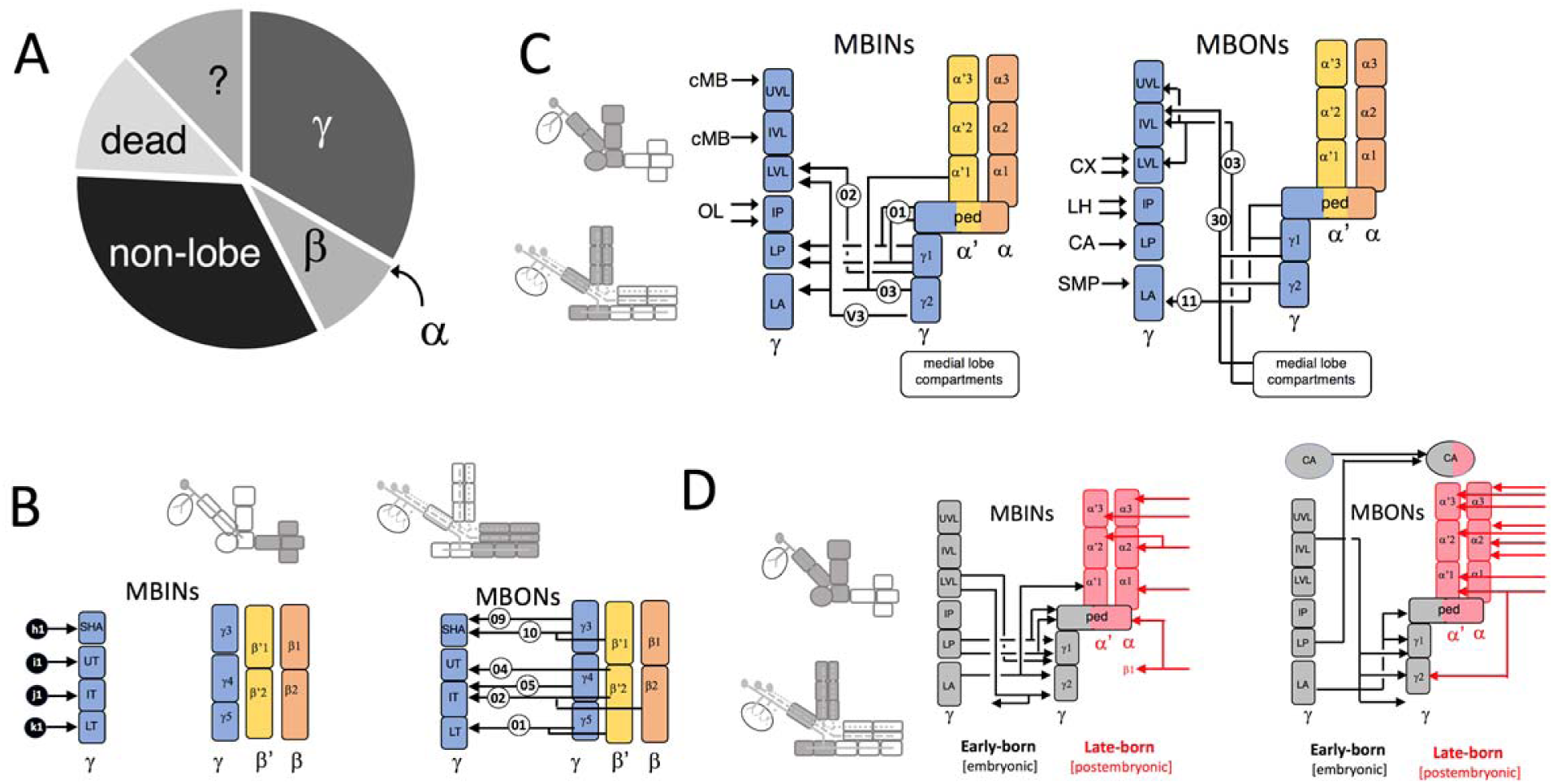
Comparison of embryonic versus postembryonic origins of the MBINs or MBONs that contribute to the mushroom body compartments of the adult. **(A)** A chart showing the percentage of larval mushroom body MBINs and MBONs that have different terminal adult fates. A few fates are unknown (?), but the remainder include death (12%), innervation of adult γ (33%), α (0%), or β lobe (9%) compartments, or non-mushroom body circuits (33%). **(B, C)** A schematic summary of the terminal fates of the neurons innervating various regions of the larval mushroom body. **(B)** the MBINs for the larval medial lobe compartments are recruited from the population of “doomed” neurons (black circles), while the MBONs serve similar roles in both the larval and adult structures. **(C)** the remaining larval compartments are supplied by neurons that assume a temporary function in the larva and then have a terminal function elsewhere in the brain, or are the larval forms of adult MBINs or MBONs (adult cell number in circles are as in Fig 2 and 3). CA: calyx; CX: central complex; cMB: circum-mushroom body neuropil; OL: optic lobe; SMP: superior medial protocerebrum. **(D)** Developmental origins of the classes of Kenyon cells that make up the core of the adult mushroom body and the MBINs and MBONs that innervate this core. The comparison is presented for the compartments of the vertical lobe, base and peduncle and is based on Figs 2, 3, and 7AB. Embryonic-born classes of Kenyon cells and their input and output cells are in black while postembryonic-born Kenyon cells and their input and output cells are in red.

Whether the association of a larval neuron with the mushroom body represents the cell’s assumed or terminal identity relates to the compartment it innervates. Seven of the larval compartments correspond to the six adult compartments that contain γ cell axons. The larval LP, LA, LVL and SHA compartments correspond to the adult peduncle, γ1, γ2 and γ3 compartments respectively. The three larval “toes” (UT, IT, and LT) correspond to compartments γ4 and γ5. The remaining 10 compartments contain axons of the α’β’ and the αβ Kenyon cells and are not directly homologous to any larval compartments because they have different Kenyon cell cores. However, the β and β’ compartments share features in common with neighboring adult γ compartments.

Figure 9B examines the extrinsic neurons of the larval medial lobe compartments from the perspective of their terminal identity. On the MBIN side, the larval compartments are innervated solely by “doomed” neurons that temporarily serve as PAM neurons but then die at metamorphosis. For the larval medial lobe MBONs, by contrast, they express their terminal identities at hatching and express the same transmitters and innervate comparable compartments along the medial lobe axis (Fig. 9B) in both larva and adult.

The metamorphic changes for the neurons in the remaining six larval compartments are variable and summarized in Figure 9C. As described above, the adult peduncle, γ1 and γ2 compartments are homologs of the larval LP, LA and LVL compartments, respectively. On the MBIN side, the three dopamine neurons to these compartments acquire their terminal identity as DANs by hatching and provide dopamine input in both larva and adult (Table 1). The octopaminergic MBIN, OAN-g1, innervates the homologous compartment in both larva and adult (LVL vrs γ2), but its terminal adult form as OA-VPM3 expands to other targets beyond the mushroom body. The only stable compartment on the output side is LA ( = γ1); in the larva MBON-d1 provides GABAergic from LA, and in its adult form of adult MBON 11, this neuron continues to provide output from the γ1 and the peduncle compartments. The larval LP and LVL compartments, by contrast, are supplied by trans-differentiated neurons. MBON-c1 provides larval-specific cholinergic output from LP and MBON-g1 & -g2 provide GABAergic output from LVL. The adult-specific MBONs that replace these cells at metamorphosis invert the transmitter output from these two compartments.

The remaining larval compartments, UVL, IVL and IP, have no adult homologs. The UVL and IVL compartments form a vertical lobe “facsimile” for the larva, thereby substituting for the lack of α and α’ axons and their extrinsic neurons. Their MBINs and MBONs assume a vertical lobe function for the larva but their terminal destinations are either in the adult medial lobes or in other parts of the adult brain (Fig.9C).

The IP compartment is intriguing because it is innervated by a unique cluster of MBINs (the PAL cluster) and it has no counterpart in the adult mushroom body. While the nine other larval compartments are highly interconnected by one- or two-step connections from the MBONs to the MBINs, the IP MBINs receive no such feedback (Eschbach *et al.,* 2020). Likewise, its MBONs provide the least amount of crosstalk to the other larval compartments. This circuit isolation suggests that the IP compartment may be involved in a type of learning distinct from that handled by other compartments. Since none of its input or output cells have terminal functions associated with the adult mushroom bodies, the type of learning mediated through the IP compartment may be unique to the larva.

Figure 9D shows that the temporal sequence of birth of the major classes of adult Kenyon cells is paralleled by the temporal sequence of birth of their respective MBONs and MBINs. We did not include the medial lobe in this comparison because of the massive postembryonic addition of the PAM neurons (Figure 7A). The γ Kenyon cells start being produced during embryogenesis and almost all their MBINs and MBONs are also produced through the same period. The latter have acquired their terminal identities by the time hatching and innervate the mushroom body in both larva and adult. By contrast, the α’β’ and αβ Kenyon cells are born late in larval life and through metamorphosis (Lee *et al*., 1999) and we find that they are supplied primarily by late-born MBONs and MBINs that arise during postembryonic growth and have no role in the larval structure.

### Changes in mushroom body circuit architecture through metamorphosis

A goal of this study was to determine the extent that mushroom body circuits were conserved through metamorphosis. Figure 10A summarizes the compartmental overlap of MBINs and MBONs in their larval versus adult configurations. There are three MBIN-MBON pairings that are found in both stages. Not unexpectedly, two involve the multi-compartmental feedback neuron, APL, which expands to cover all of the compartments in the adult. The last conserved pairing occurs outside of the lobes, in the calyx. For the uni-compartmental neurons of the lobe system, though, we found no MBIN-MBON pairings that persisted through metamorphosis.

**Figure 10.**
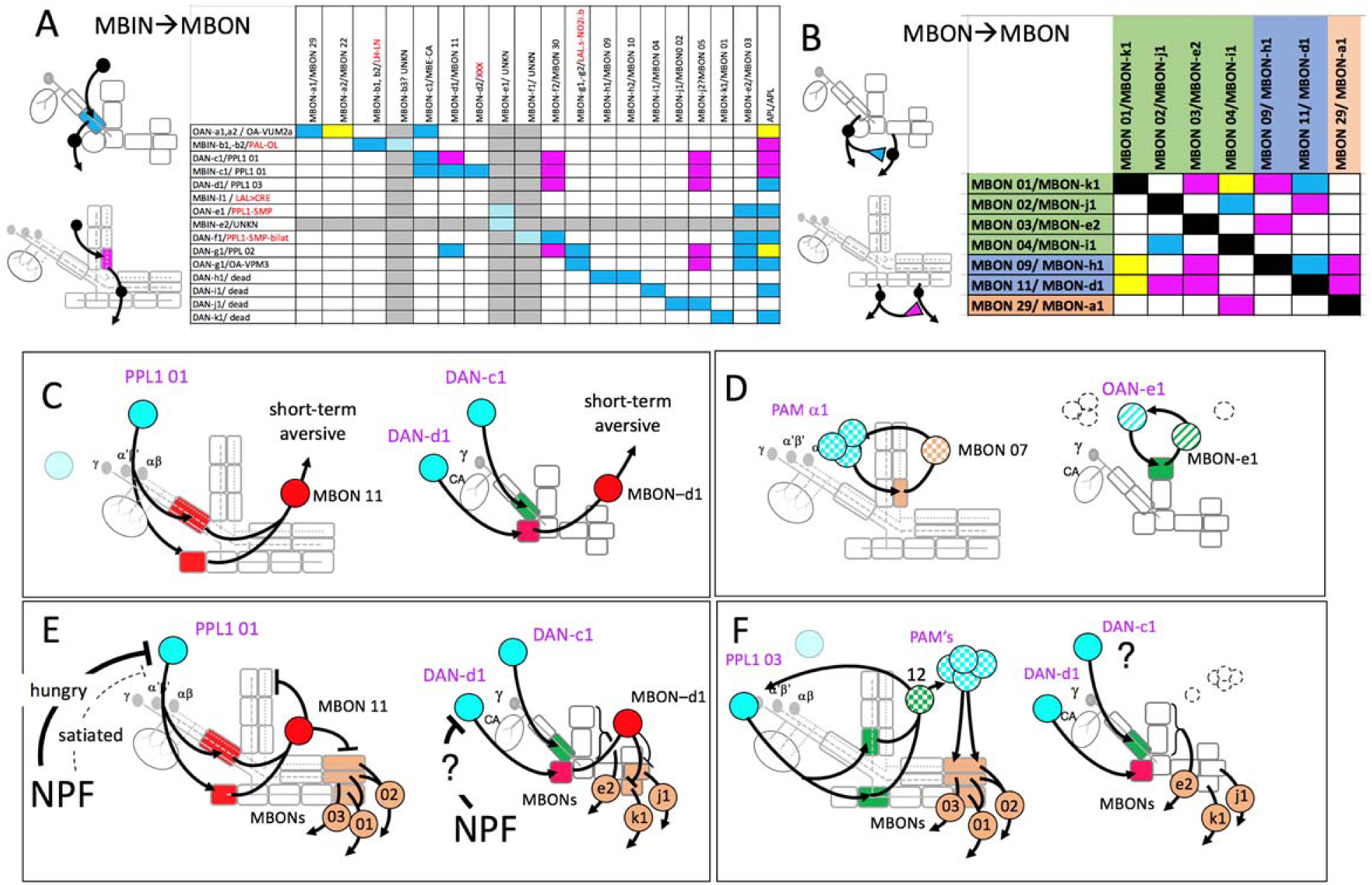
Fate of circuit connections in the mushroom bodies through metamorphosis. **(A)** Matrix showing the overlap of MBIN axon terminal with MBON dendritic trees in the same compartment. Blue: larval pairings; red: adult pairings; yellow: found in both. Cells and columns that are grayed out are ones for which the adult identity is unknown. Larval/adult names are provided for each cell, with the red names being the terminal identity of neurons that leave the larval mushroom bodies. **(B)** Matrix showing the larval (blue) and adult (red) connectivity for a set of MBONs that maintain this function through metamorphosis. **(C-F)** comparison of the larval and adult states of circuit components for specific examples of adult circuits that illustrate various degrees of learning complexity: **(C)** Simple short-term aversive conditioning. **(D)** Consolidation of appetitive conditioning. **(E)** Shift the valence of a learned response in response to change in internal state (hunger). **(F)** Re-consolidation and extinction of memories based on subsequent experience. Extrinsic neurons with mushroom body function only in the larva are cross-hatched and those with adult-specific mushroom body function are stippled in the adult and shown as dashed outlines in the larva. Cell body and compartment colors: cyan, aminergic; red: GABAergic; green: cholinergic; tan: glutaminergic. See text for details.

Besides MBIN to MBON connections, the compartments of both the larva (Eichler *et al*., 2017; Eschbach *et al*., 2020) and the adult (Aso *et al*., 2014; Li *et al*., 2020) are highly interconnected, both by MBON to MBON connections and by feedback and feed forward connections from MBONs back to MBINs. For MBON to MBON interactions, larval (Eichler *et al*., 2017) and adult (Li *et al*., 2020) data are available for seven of the MBONs that function in both circuits (Fig 10B). There are 42 possible pair-wise connections amongst these cells excluding connections of a MBON onto itself. The MBONs are more highly interconnected in their adult configuration as compared to their larval one: the adult group shows 13 connections (31% of possible connections) while in their larval configuration they have seven (17%). Importantly, only three connections (7%) are common to both configurations. This percentage is similar to the 5% predicted if the two stages were wired up completely independently at their respective levels. This low level of shared connections suggests that in both configurations the MBONs are free to interconnect in a way that is optimal for the particular life stage.

There are very few direct MBON to MBIN connections in the larva. Rather, connections between these cells are provided by an extensive network of one- and two-step feedback and feed forward pathways (Eschbach *et al*., 2020). We do not know the metamorphic fates of the neurons of these pathways, so we cannot compare their connectivity in the two stages.

The simplest functions of the mushroom bodies are in short-term appetitive or aversive conditioning and there are examples in both larvae and adults showing that manipulation of the input or output cell from a single compartment can support or suppress these short-term processes (*e.g.,* Saumweber *et al*., 2018; Aso *et al*., 2012). Although their input-output relationships do not survive metamorphosis (Figure 10A), single neurons may still have similar functional roles in both larva and adult, such as diagramed in Figure 10C. In the adult, the stimulation of the DAN PPL1 01 (= PPL1-γ1pedc ) is sufficient to induce short-term aversive conditioning to a paired odor (Aso *et al.,* 2012; Das *et al.,* 2014). This neuron innervates the peduncle and γ1 compartments and acts through the GABAergic output cell MBON-11 (Aso *et al.,* 2012; Aso *et al.,* 2014). The larval form of PPL1 01 is DAN-c1 (Figure 2), but it only targets the larval LP compartment where it contacts the cholinergic output neuron MBON-c1 (Eichler *et al*., 2017). Stimulation of DAN-c1 in the larva, though, does not support short-term aversive conditioning (Eschbach *et al*., 2020). The larval version of the adult output partner (MBON 11) is called MBON-d1 (Figure 3). It receives input from DAN-d1 (Eichler *et al.,* 2017), and stimulation of this DAN-d1 is sufficient to support short-term aversive conditioning in the larva (Eschbach *et al.,* 2020). Consequently, MBON-d1/ MBON 11 is involved in short-term aversive conditioning in both larva and adult, but a different dopamine MBIN instructs it in the two stages.

Mushroom bodies of larvae and adult also mediate higher order functions such as long-term learning, shifts in learning valence dependent on internal state, and the ability to extinguish or re-consolidate memories (Cognigi *et al.,* 2018; Thum & Gerber, 2019). These higher order processes involve intra- and intercompartmental connections with feed-forward or feed-back interactions amongst MBONs or from the MBONs back to the MBINs. Figures 10D-F illustrate the circuitry underlying three types of higher order networks in the adult and indicate which components are present in the larva. Memory consolidation continues for minutes to hours after training and involves recurrent activity within α lobe compartments. A recurrent loop that is necessary for memory stabilization after training with a sugar reward involves PAM-α1 neurons, α Kenyon cells and the cholinergic MBON-α1’s (Ichinose *et al*., 2015). Larvae lack all three of these neuron types, but Eichler *et al.,* (2015) showed that the larval vertical lobe facsimile has similar circuit motifs that involve feedback of cholinergic MBONs back onto their compartmental MBINs, such as that involving OAN-e1 and MBON-e1 (Figure 10D). The latter loop potentially provides the larva with the recurrent circuits that might consolidate memories, but these are replaced by the postembryonic-born neurons that provide this function in the adult.

Figure 10E shows an example of a feed-forward pathway that allows the valence of a learned response to change dependent on a fly’s internal state. Beyond their involvement in short-term aversive conditioning, the adult PPL1 01/MBON-11 pair functions to adjust how flies respond to a learned, aversive odor based on their hunger state (Perisse *et al.,* 2016). Amongst its other targets, MBON 11 inhibits three medial lobe MBONs (Oswald *et al.,* 2015). The activity of the latter MBONs promotes avoidance behavior while their suppression promotes approach (Oswald et al., 2015; Perisse *et al.,* 2016). Neuropeptide F (dNFP) induces a hunger state in the fly; its release is elevated in hungry flies and suppressed in satiated flies. In the absence of dNPF in the satiated fly, PPL1 01 activity suppresses the inhibitory MBON 11, thereby derepressing the medial lobe MBONs and promoting avoidance behavior. In the hungry fly, dNPF release suppresses PPL1 01, resulting in elevated MBON-11 activity which suppresses the medial lobe MBONs and reduces avoidance. dNPF also enhances sugar learning in larvae (Rohwedder *et al.,* 2015). The neurons involved in the adult circuit are also present in the larva, but MBON-d1, the larval form of MBON 11, has a reduced set of MBON targets; it has moderate connections to MBON-k1 ( = adult MBON 01), a weak connection to MBON-j1 ( = adult MBON 02), and no connection to MBON-e2 ( = adult MBON 03) (Eichler *et al.,* 2017). A more important difference from the adult circuit is that MBON-d1 is instructed by a different DAN (DAN-d1) in the larva versus the adult as described above. It is not known if DAN-d1 is a larval target for dNPF and, thereby, provides an analogous pathway for hunger state to modify learning in the larva.

Figure 10F shows an adult circuit involved in memory re-consolidation and extinction (see Cognigi *et al.,* 2018). PPL1 03 (γ2α’1) activates the cholinergic MBON 12 (MBON-γ2α’1) which provides feed-back excitation to PPL1 03 and also feeds across to PAM DANs that innervate the three medial lobe MBONs described above. The larva possesses the first and last cells in this circuit but lacks both the PAMs and the critical MBON 12. In its larval state, we find that PPL1 03 is DAN-d1. DAN-d1 innervates the LA compartment which has a GABAergic output through MBON-d1. This inhibitory output is ill adapted for feed-back and feed-across excitation in the larva. It is possible that this function in the larva has shifted to DAN-c1 and is associated with switch in cholinergic output from the LP compartment of the larva to the γ2 and α1 compartments of the adult. There are no direct connections from MBON-c1 back to DAN-c1 (Eichler *et al.,* 2017), but the feed-forward, feed-back, and feed-across connections described by Eschbach *et al*. (2020) may provide the required pathways.

### The persistence of memory traces through metamorphosis

Experiments on aversive conditioning of *Drosophila* larvae suggested that the memory of larval training can endure through metamorphosis (Tully *et al.,* 1994). Our anatomical analysis did not identify any circuit elements that may support the persistence of a memory trace from larva to adult. Within the lobe system, none of the MBIN-MBON pairings persist (Figure 10A) and persisting MBON to MBON connections are rare (Figure 10B). A memory trace might involve more complex pathways as described by Eschbach *et al.,* 2020, but these cannot be addressed in this study.

Our failure to find anatomical support for persistence of a memory trace from larva to adult in *Drosophila* should not be generalized to other insects that undergo complete metamorphosis. There is compelling evidence that learning in caterpillars and beetle grubs can carry through to the adult (Blackistin *et al.,* 2008, 2015). Butterfly and beetle larvae have an extended embryonic development and hatch with a more complex nervous systems as compared to *Drosophila* larvae. The extended period of embryonic neurogenesis means that caterpillars and grubs hatch with more of the neuron types needed for constructing a functional mushroom body and, therefore, are less dependent on using trans-differentiation to generate missing neuron types. A higher incidence of the same neurons being used in the mushroom body circuits of both stages increases the likelihood that some connections persist through metamorphosis.

### The evolution of a larval mushroom body

The insect metamorphic life history, which involves making different larval and adult nervous systems, arose from an ancestral condition in which the embryo generated a single nervous system that served both the nymph and the adult. In the latter, even neurons that have adult-specific functions such as those involved in flight or reproduction already have their terminal form in the hatchling nymph (reviewed in Truman, 2005). Our study of the origins and fates of mushroom body neurons in *Drosophila* provides insight into how this second nervous system may have been intercalated into an ancestral plan that generated only one CNS (Figure 11). Other strategies are evident in the formation of the central complex in larval beetles (Farnworth *et al*., 2020).

**Figure 11.**
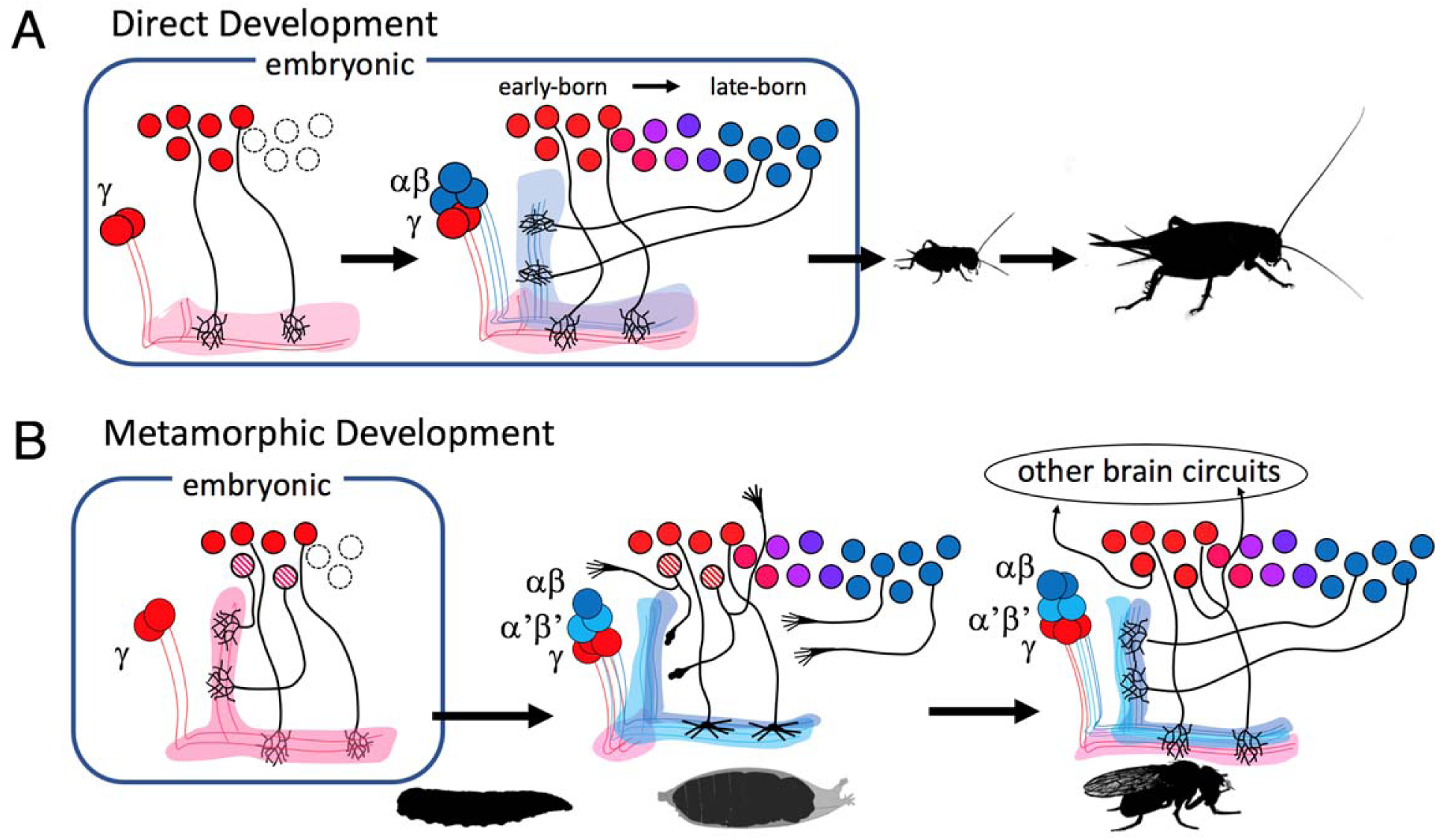
The proposed scheme showing the relationship of the development of a single mushroom body in a direct developing insect to the production of two sequential versions of the mushroom body in a metamorphic insect like *Drosophila*. **(A)** In the direct developing insects γ Kenyon cells are made before the αβ type Kenyon cells. The γ class generates a medial lobe while the αβ classes provide a vertical lobe(s) as well as more medial lobe axons. The timing of birth of the input and output cells to the Kenyon cell classes is assumed to parallel the relative time of birth of their targets with those innervating γ neurons being born before those innervating the αβ neurons. **(B)** In the derived, metamorphic system, typified by *Drosophila,* the early hatching of the larva results in only γ neurons being present, and these possess a novel, vertical axon to make up for the absence of the αβ neurons. The input and output cells to the γ neurons are also present at hatching and some of these assume their expected roles along the larval medial lobe. The larval vertical lobe, though, is analogous, but not homologous, to the adult structure. Its core is composed of γ cell axons, rather than α branch axons, and none of the later-born neurons that innervate α axons are present at hatching. The role of the latter neurons is taken over by early-born neurons that trans-differentiate to assume mushroom body functions in the larva but then revert to their ancestral functions in other brain circuits at metamorphosis. By this latter time the late-born classes of Kenyon cells and their input and output cells have been born and an adult mushroom body is formed using the same cell classes as used in their direct developing relatives.

Direct developing insects like the house cricket, *Acheta domesticus*, produce two classes of Kenyon cell classes, the γ and the αβ classes. Both are produced during embryogenesis with the γ set being made before the αβ cells (Malaterre *et al*., 2002). With the exception of the optic lobes (Anderson, 1978) and the addition of more αβ Kenyon cells (Malaterre *et al*., 2002), direct developing insects finish producing neurons for their CNS during embryogenesis (Shepherd & Bate, 1990; Truman & Ball, 1998). Consequently, all of their mushroom body extrinsic neurons should be present at hatching, and we assume that the relative times of their birth mirrors that of their Kenyon cell targets. In other words, we assume that the MBONs and MBINs innervating the γ cells are born before those innervating the αβ neurons, and that both sets are present in the first stage nymph. Unfortunately, though, the temporal sequence by which these cells arise has not yet been determined for any direct developing insect.

Compared to their direct developing relatives, *Drosophila* embryos undergo an early arrest in neurogenesis, resulting in a larval brain that contains about 10% of the neurons found in the adult central brain. We find that this truncation results in the presence of γ neurons and many of the early-born extrinsic neurons that innervate them, but the later-born αβ Kenyon cells and their input and output neurons are absent. The terminal fates of these earliest-born neurons are for the medial lobe and many are used as such for the larva, but making a vertical lobe is a problem because of the lack of αβ and α’β’ Kenyon cells and their extrinsic neurons. We find that the larva solves this problem by constructing a “ facsimile” of a vertical lobe. This involves modifying the γ Kenyon cells with a larval-specific vertical axon branch to form the core of the vertical lobe facsimile and finding substitutes for the missing vertical lobe MBINs and MBONs. The latter were recruited from early-born neurons that were destined for adult circuits not needed in the larva or from “spare” medial lobe MBONs that are redirected to the vertical lobe. These neurons assume a temporary identity that is maintained through larval life but, when the larval vertical lobe facsimile is disassembled at metamorphosis, they trans-differentiate to assume their terminal identity in the adult brain. As the larval system is being disassembled, the αβ and α’β’ neurons and their late-born extrinsic cells then construct the adult vertical lobes. Consequently, from the perspective of the adult, the relative temporal ordering of the production of the Kenyon cell types and their corresponding MBINs and MBONs remains the same as is thought to exist in the direct developing insects from which they evolved.

The ability of some neurons to trans-differentiate and thereby perform different functions in the larva versus the adult seems crucial for making a sophisticated larval CNS. Our observations on the metamorphic fates of the lineage 13B leg interneurons (Figure 6) argue that the adult identities of these neurons are their ancestral identities, and their assumed, larval identities are a derived condition supporting the evolution of a larva. A seminal paper by Thomas *et al*. (1984) indicated that the neuron types for the insect CNS are generated according to a set of highly conserved spatial and temporal rules that have changed little through the span of insect evolution. Our findings for the adult mushroom bodies are consistent with this notion, but the production of some cell types for the larval system, such as the early appearance of MBINs and MBONs for the larva vertical lobe, appear to break these rules, While the assumed identity and function of these neurons in the larva appear to deviate from the ancestral plan for making neuron types, we find that their terminal identities are in accord with it. In other words, the ability of some neurons to trans-differentiate allows them to both maintain the rules for generating neuronal phenotypes for the adult while also temporarily suspending or modifying such rules to generate an assumed identity for these cells while in the larva. Trans-differentiation allowed these neurons to uncouple their larval functions from those of the adult, thereby allowing selection to potentially modify one version of the cell without impacting the other. The evolutionary success of insects with complete metamorphosis is attributed to the larval and adult stages being based on independent developmental modules that allow selection to change one without compromising the other [Yang, 2001]. Trans-differentiating neurons provide a cellular example of such an uncoupling. We do not yet know how these neurons achieve an uncoupling that allows them to assume two different identities during the life of the animal, although, as speculated above, we expect that temporary changes may occur within lineages to alter the temporal or spatial information that direct neuronal identity. Besides providing insight into a key innovation in insect evolution, understanding natural mechanisms by which cell fates are changed in insects may reveal new mechanisms that will be useful in changing neuronal phenotypes in other animals to deal with issues in aging or disease.

## Materials & Methods

### Fly stocks

Drosophila stocks were raised on standard corn meal-molasses at either 25°C or room temperature. The genetic stocks used in this study are summarized in Tables 4 and 5.

**Table 4.** Lines used to determine fates of larval neurons.

| Cell name | split line | split line | split line |
| --- | --- | --- | --- |
| MBIN-b1,b2 | **SS21716 (FS, TL)** |  |  |
| DAN-c1 | **SS03066 (FS, TL)** | **MB586B (FS)** | SS01702 (FS) |
| DAN-d1 | **MB328B (FS, TL)** |  |  |
| MBIN-l1 | **SS04484 (FS, TL)** | SS01624 (FS) |  |
| OAN-e1 | **SS36923 (FS)** | SS01958 (FS) |  |
| DAN-f1 (+ DAN-c1) | **MB065b (FS, TL)** | MB145 (TL) |  |
| DAN-g1 | **SS01716 (FS, TL)** | SS01755 (FS) |  |
| OAN-g1 (sVPMmx) | **SS25844 (FS)** | **SS04268 (FS)** |  |
| DAN-h1 | **SS01949 (NC,TL)** | SS01696 (NC) | MB440B (NC) |
| DAN-i1 | **SS01949 (NC,TL)** | MB196C (NC) |  |
| DAN-j1 | **SS01949 (NC,TL)** | MB316B (NC) | MB340C (NC) |
| DAN-k1 | **SS01757 (NC,TL)** | MB198B (NC) | SS00616 (NC) |
| MBON-a1 | **SS00867 (FS,TL)** | **SS01417 (FS)** |  |
| MBON-a2 | **SS02006 (FS)** |  |  |
| MBON-b1,b2 | **SS01708 (FS, TL)** | **SS04112 (FS)** | SS01959 (FS) |
| MBON-c1 | SS21789 (FS, TL) |  |  |
| MBON-d1 | **SS01705 (FS,TL)** |  |  |
| MBON-d2 | **SS04231 (FS)** |  |  |
| MBON-e2 | SS04559 (FS, TL) |  |  |
| MBON-f2 | **SS04328 (FS, TL)** | SS36248 (FS) |  |
| MBON-g1,g2 | **SS02130 (FS, TL)** | SS02121 (FS) |  |
| MBON-h1,h2 | **SS01725 (FS, TL)** |  |  |
| MBON-i1 | SS01771 |  |  |
| MBON-j1 | **SS01973 (FS,TL)** | SS01972 (FS) |  |
| MBON-j2 | **SS00860 (FS,TL)** |  |  |
| MBON-k1 | **SS01962 (FS)** | **SS01980 (TL)** |  |
| APL | **SS01671 (FS, TL)** |  |  |

FS: flip-switch immortalization; NC: no adult counterpart; TL: developmental timeline. **Bold** lines were the best lines for each cell.

**Table 5.** Split GAL4 lines used in study.

| split line | target cell | AD | DBD |
| --- | --- | --- | --- |
| **MB065b** | DAN-f1 (+ DAN-c1) | TH-p65ADZp in attP40 | R72B05-ZpGdbd in attP2 |
| MB145 | DAN-f1 (+ DAN-c1) | R15B01-p65ADZp in attP40 | R72B05-ZpGdbd in attP2 |
| MB196C | DAN-i1 | R58E02-p65ADZp in attP40 | R36B06-ZpGdbd in attP2 |
| MB198B | DAN-k1 | R58E02-p65ADZp in attP40 | R71D01-ZpGdbd in attP2 |
| MB316B | DAN-j1 | R58E02-p65ADZp in attP40 | R93G08-ZpGdbd in attP2 |
| **MB328B** | DAN-d1 | R82C10-p65ADZp in attP40 | R32F01-ZpGdbd in attP2 |
| MB340C | DAN-j1 | R93D10-p65ADZp in attP40 | R12G04-ZpGdbd in attP2 |
| MB440B | DAN-h1 | R30G08-p65ADZp in attP40 | R17D06-ZpGdbd in attP2 |
| **MB586B** | DAN-c1 | TH-p65ADZp in attP40 | R72G06-ZpGdbd in attP2 |
| SS00616 | DAN-k1 | 71D01-p65ADZp in VK00027 | 17D06-ZpGdbd in attP2 |
| **SS00860** | MBON-j2 | w; R89G07-p65ADZ; MKRS/TM6B | R24E12-ZpGdbd in attP2 |
| **SS00867** | MBON-a1 | w; R93G12-p65ADZ; MKRS/TM6B | R52E12-ZpGdbd in attP2 |
| **SS01417** | MBON-a1 | w; R52E12-p65ADZp | R93G12-ZpGdbd in attP2 |
| SS01624 | MBIN-l1 | w; R84D07-p65ADZ | R37G09-ZpGdbd in attP2 |
| **SS01671** | APL | R21D02-p65ADZp | R55D08-ZpGdbd in attP2 |
| SS01696 | DAN-h1 | 76F05-p65ADZp in attP40 | 95H02-ZpGdbd in attP2 |
| SS01702 | DAN-c1 | VT054895-p65ADZ in attP40 | R53C05-ZpGdbd in attP2 |
| **SS01705** | MBON-d1 | R11E07-p65ADZp in attP40 | R52H01-ZpGdbd in attP2 |
| **SS01708** | MBON-b1,b2 | R12G03-p65ADZp in attP40 | 21D02-ZpGdbd in attP2 |
| **SS01716** | DAN-g1 | R14E06-p65ADZp in attP40 | R27G01-ZpGdbd in attP2 |
| **SS01725** | MBON-h1,h2 | R20A02-p65ADZp in attP40; MKRS/TM6B | R28A10-ZpGdbd in attP2 |
| SS01755 | DAN-g1 | R46F09-p65ADZp | R14E06-ZpGdbd in attP2 |
| **SS01757** | DAN-k1 | w; R48F09-p65ADZp; MKRS/ TM6B | R27A11-ZpGdbd in attP2 |
| SS01771 | MBON-i1 | w; 65A05-p65ADZ; MKRS/TM6B | 14C08-ZpGdbd in attP2 |
| **SS01949** | DAN-h1, -i1, -j1 | VT026700-p65ADZp in attP40 | VT058464-ZpGDBD in attP2 |
| SS01958 | OAN-e1 | VT023826-p65ADZp in attP40 | R75F01-ZpGdbd in attP2 |
| SS01959 | MBON-b1,b2 | VT027952-p65ADZp in attP40 | R26A02-ZpGdbd in attP2 |
| **SS01962** | MBON-k1 | VT033301-p65ADZp in attP40 | R27G01-ZpGdbd in attP2 |
| SS01972 | MBON-j1 | VT057469-p65ADZp in attP40 | 12C11-ZpGdbd/ TM3 in attP2 |
| **SS01973** | MBON-j1 | VT057469-p65ADZp in attP40 | R18D09-ZpGdbd in attP2 |
| **SS01980** | MBON-k1 | VT020613-p65ADZp in attP40 | VT033301-ZpGdbd in attP2 |
| **SS02006** | MBON-a2 | w; 93G12-p65ADZ; MKRS/TM6B | 71E06-ZpGdbd in attP2 |
| SS02121 | MBON-g1,g2 | R21D06-p65ADZp in attP40 | R23B09-ZpGdbd in attP2 |
| **SS02130** | MBON-g1,g2 | w; R23B09-p65ADZp; MKRS/ TM6B | R21D06-ZpGdbd in attP2 |
| **SS03066** | DAN-c1 | VT054895-p65ADZ in attP40 | VT057278-ZpGdbd in attP2 |
| **SS04112** | MBON-b1,b2 | VT027952-p65ADZp in attP40 | HAV5; CyO/Sco; 21D02-ZpGDBD in attP2 |
| **SS04231** | MBON-d2 | VT032899-p65ADZp in attP40 | HAV5; CyO/Sp; 87G02-ZpGDBD in attP2 |
| **SS04268** | OAN-g1 (sVPMmx) | VT012639-p65ADZp in attP40 | VT016127-ZpGdbd in attP2 |
| **SS04328** | MBON-f2 | VT033301-p65ADZp in attP40 | VT029593-ZpGdbd in attP2 |
| **SS04484** | MBIN-l1 | R37G09-p65ADZp in attP40 | VT007174-ZpGdbd in attP2 |
| **SS21716** | MBIN-b1,b2 | VT048835-p65ADZp in attP40 | VT026664-ZpGdbd in attP2 |
| SS21789 | MBON-c1 | VT050247-p65ADZp in attP40 | VT050247-ZpGDBD in attP2 |
| **SS25844** | OAN-g1 (sVPMmx) | VT040569-p65ADZp in attP40 | VT061921-ZpGdbd in attP2 |
| SS36248 | MBON-f2 | VT016795-p65ADZ in attP40 | VT029593-ZpGdbd in attP2 |
| **SS36923** | OAN-e1 | VT054895-p65ADZ in attP40 | HAV5; CyO/Sco; 75F01-ZpGDBD in attP2 |
| SS04559 | MBON-e2 | w; 65A05-p65ADZ; MKRS/TM6B | VT045663-ZpGDBD in attP2 |

### Flp-Switch treatments

The expression pattern seen in the late 3^rd^ instar in stable spilt lines was maintained through metamorphosis using the Flip-Switch method described in Harris *et al*. (2015). Using a similar strategy of the gene-switch method (Roman *et al.,* 2001), flippase was fused to the ligand-binding domain of the human progesterone receptor, rendering it dependent on progesterone or a progesterone mimic to move into the nucleus. Stable spilt lines were crossed to *pJFRC48-13XLexAop2-IVS-myrtdTomato in su(Hw)attP8; Actin5Cp4.6>dsFRT>LexAp65 in su(Hw)attP5; pJFRC108-20XUAS-IVS-hPRFlp-p10 in VK00005/TM6*.

We used the progesterone mimic mifepristone (RU486, Sigma Aldrich; #M8046) to cause translocation of the flippase to the nucleus where it could then flip-out the STOP cassette in the Actin-LexAp65 transgene. We used surface application of RU486 to food vials. Parents were allowed to lay eggs in a food vial for a few days, then transferred to a fresh vial.

Approximately 4 days after transfer, 60μl of a ∼10mM RU486 stock solution (10 mg RU486 dissolved in 2 ml 95% ethanol) was applied to the surface of the food. At 24 h after treatment, any larvae that had wandered and/or pupariated were discarded to ensure that test animals had fed on RU486 for at least 24 h. At 48h after treatment, the subsequent wandering larvae and pupae (which had all fed on RU486 for 24-48 h during the L3 stage) were collected and transferred to an untreated food vial. These animals were then dissected in Schneider’s S2 culture medium as adults. This treatment results in constitutive LexA expression in any cells that express GAL4 during the L3 stage, but, because the RU486 persists at least partway through metamorphosis neurons that start expressing in early to mid-metamorphosis also show up.

### Lineage-targeted twin-spot MARCM

Specific neuronal lineages were targeted using lineage-restricted drivers (Awasaki *et al*., 2014) to label sister clones with twin-spot MARCM (Yu *et al*., 2009). The Vnd-GAL4 driver permits targeting 18 fly central brain lineages, including the FLAa2 lineage (Lee *et al*., 2020); and stg14-GAL4 driver covers eight type II neuronal lineages, including the DL1 lineage (Wang *et al*., 2014). Twin-spot clones were induced at specific times after larval hatching and examined at the adult stage by immunostaining and confocal imagining, following published work (e.g. Yu *et al*., 2010).

### Preparation and examination of tissues

Tissues were dissected in PBS (phosphate-buffered saline, pH 7.8) and fixed in 4% buffered formaldehyde overnight at 4°C. Fixed tissues were rinsed in PBS-TX (PBS with 1% Triton X-100, Sigma), then incubated overnight at 4°C in a cocktail of 10% normal donkey serum (Jackson ImmunoResearch), 1:1000 rabbit anti-GFP (Jackson ImmunoResearch), 1:40 rat anti-N-Cadherin (Developmental Studies Hybridoma Bank), and 1:40 mouse anti-Neuroglian or a 1:200 dilution of mouse anti-FasII (both Developmental Studies Hybridoma Bank). For visualization of tdTomato, a 1:500 dilution of rabbit anti-DsRed (CloneTech) was substituted for the anti-GFP and the anti-Neuroglian was omitted. After repeated rinses PBS-TX, tissues stained for GFP were incubated overnight at 4°C with 1:500 AlexaFluor 488-conjugated donkey anti-rabbit, AlexaFluor 594-conjugated donkey anti-mouse, and AlexaFluor 649-conjugated donkey anti-rat (all from Invitrogen). For visualization of RFP staining was with a 1:500 dilution of AlexaFluor 594-conjugated donkey anti-rabbit and AlexaFluor 649-conjugated donkey anti-rat. After exposure to secondaries, tissues were then washed in PBS-TX, mounted onto poly-Lysine-coated coverslips, dehydrated through an ethanol series, cleared in xylenes, and mounted in DPX mountant (Sigma-Aldrich). Nervous systems were imaged on a Zeiss LSM 510 confocal microscope at 40x with optical sections taken at 2μm intervals. LSM files were contrast-enhanced as necessary and *z-*projected using ImageJ (http://rsbweb.nih.gov/ij/). Reagents summarized in Table 6.

**Table 6:** Reagents used in the present study.

| **Reagent** | **Source** | **Catalog number** |
| --- | --- | --- |
| Mouse anti-bruchpilot | Developmental Studies Hybridoma Bank | nc82-s |
| Rat anti-N cadherin | Developmental Studies Hybridoma Bank | DN-Ex #8 |
| Mouse anti-neuroglian | Developmental Studies Hybridoma Bank | BP 104 |
| Mouse anti-Fasciclin II | Developmental Studies Hybridoma Bank | 1D4 |
| Rabbit anti-DsRed | ClonTech | #632496 |
| Normal Donkey Serum | Jackson ImmunoResearch | #017-000-121 |
| AF488 Donkey α-rabbit | Jackson ImmunoResearch | #711-545-152 |
| AF488 Donkey α-Mouse | Jackson ImmunoResearch | #711-585-151 |
| AF594 Donkey α-Rabbit | Jackson ImmunoResearch | #711-585-152 |
| AF594 Donkey α-Mouse | Jackson ImmunoResearch | #715-585-151 |
| AF649 Donkey α-Rat | Jackson ImmunoResearch | #712-605-153 |
| Mifepristone (RU-486), $\geq 98\%$. | Sigma Aldrich | #M8046-100MG |
| S2 – Schneider's Insect Medium | Sigma Aldrich | #S01416 |
| DPX mountant | Electron Microscopy Sciences | # 13512 |

## Acknowledgements

We are grateful to Scarlett Pitts and Todd Laverty of the Janelia FlyCore for dealing with *Drosophila* maintenance and setting up the needed crosses. Members of the Janelia FlyLight team including Geoffrey Meissner, Susana Tae, Jennifer Jeter, Scott Miller and Sophia Protopapas were involved in dissection, immunocytochemistry, and imaging. We think Lynn Riddiford for critical comments on the manuscript. The research was funded by HHMI.

## Competing Interests

The authors have no competing interests.

## Supplemental Figures

**Figure 2---figure supplement 1.**
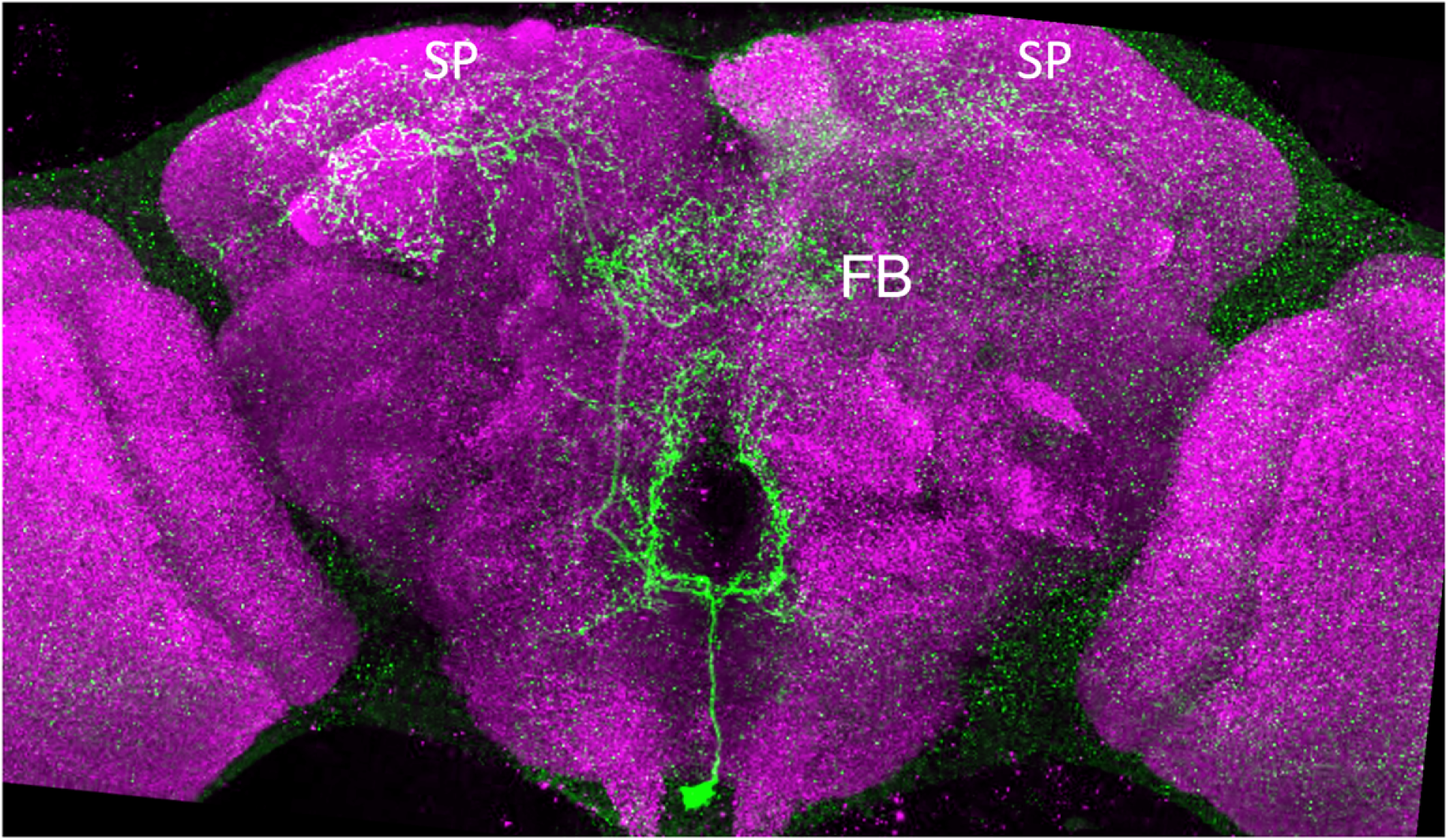
Confocal projections showing the terminal, adult structure of the larval neuron OAN-g1. The adult cell is called OA-VPM3. FB: fan shaped body; SP: superior protocerebrum. Green: pseudo color representation of RFP; Magenta: nc82.

**Figure 2---figure supplement 2.**
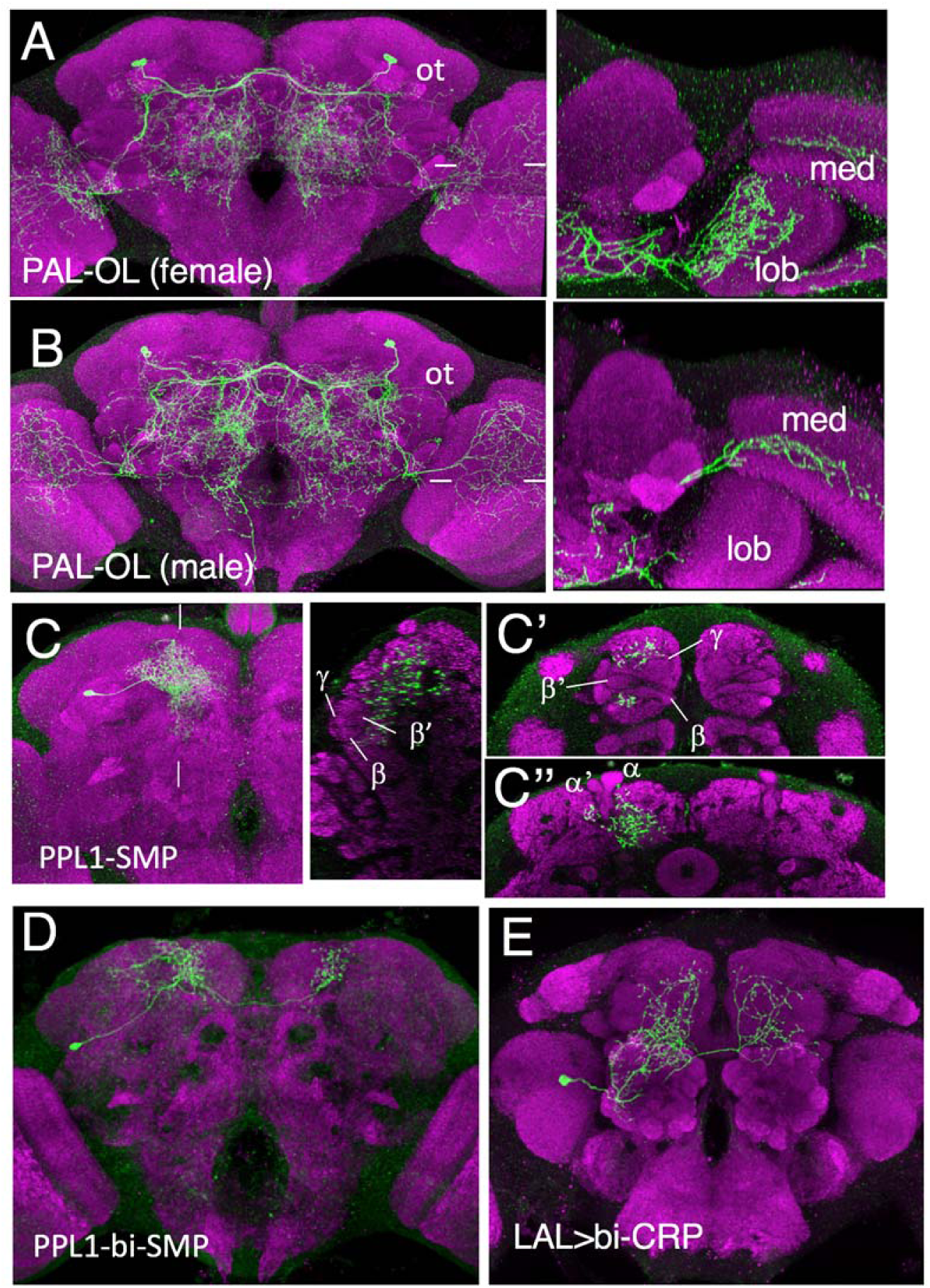
Confocal projections showing the terminal, adult identity of larval MBINs that undergo trans-differentiation at metamorphosis. **(A,B)** female and male versions of larval MBIN-b1 &-b2. Images to the right are a horizontal section at the level of the tick marks showing that in males the cell innervates the medulla (med) but in the female the medulla projection is reduced but it has extensive branching in the lobula (lob). Ot: optic tubercle. **(C)** Frontal projection showing the terminal adult morphology of larval cell OAN-e1. Image to right is a lateral section at level of the tick marks showing that the arbor is outside of the bundles of Kenyon cell axons. C’ and C” are frontal slices at levels to relationship of arbor th the γ, β’ and β lobes of the medial loves (C’) and the α and α’ lobes of the vertical lobes (C”). **(D)** terminal adult anatomy of larval DAN-f1. **(E)** Terminal adult identity of larval MBIN-l1. Green: pseudo color representation of RFP; Magenta: nc82.

**Figure 3---figure supplement 1.**
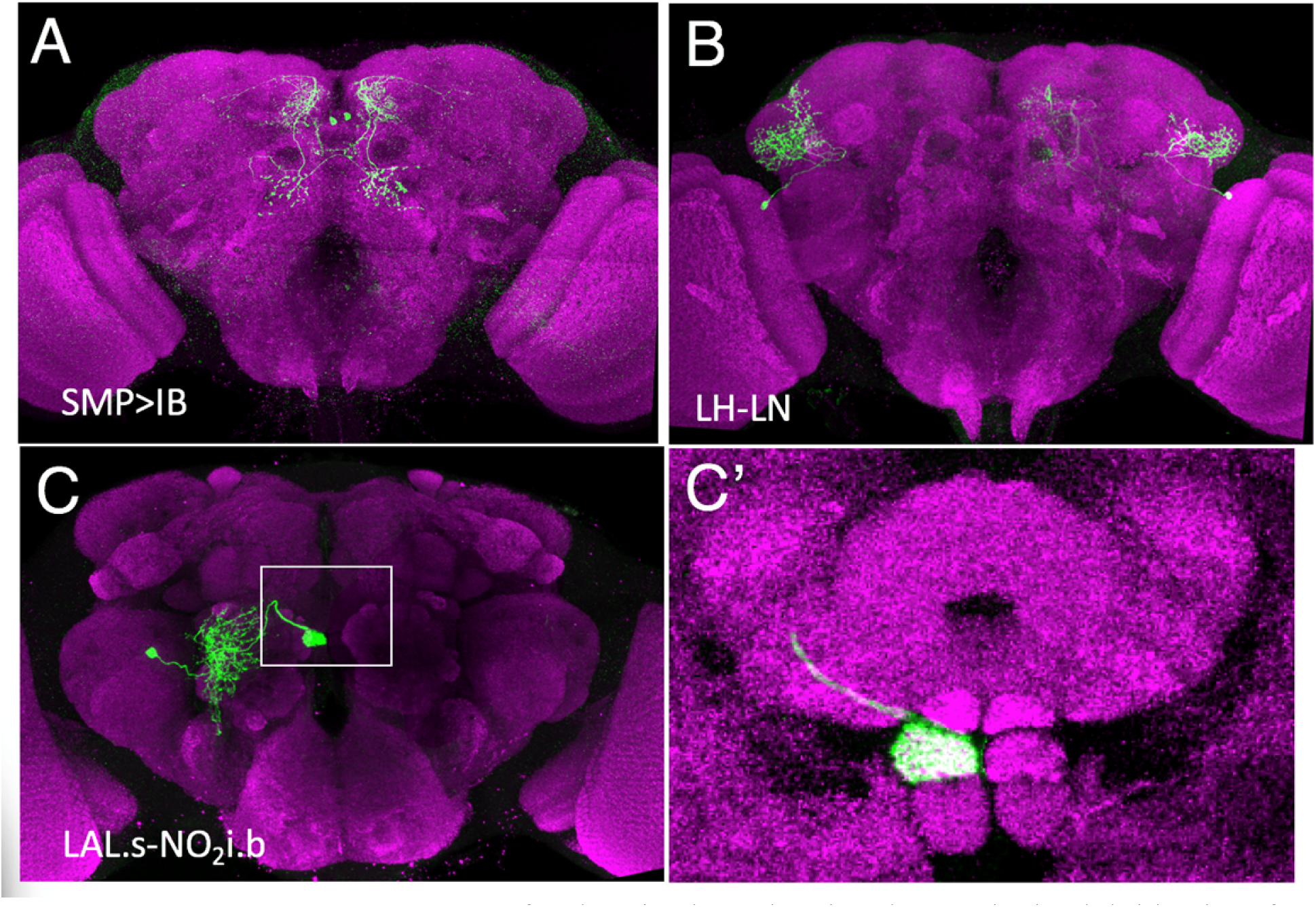
Confocal projections showing the terminal, adult identity of larval MBONs that undergo trans-differentiation at metamorphosis. Frontal views of the adult brain showing the terminal identity of **(A)** MBON-d2, **(B)** MBON-b1 and -b2, **(C)** MBON-g1 and g2. **(C’)** a magnified image of the boxed region of “C” showing the terminals of the neuron in the intermediate section of the nodulus. Green: pseudo color representation of RFP; Magenta: nc82.

**Figure 4---figure supplement 1.**
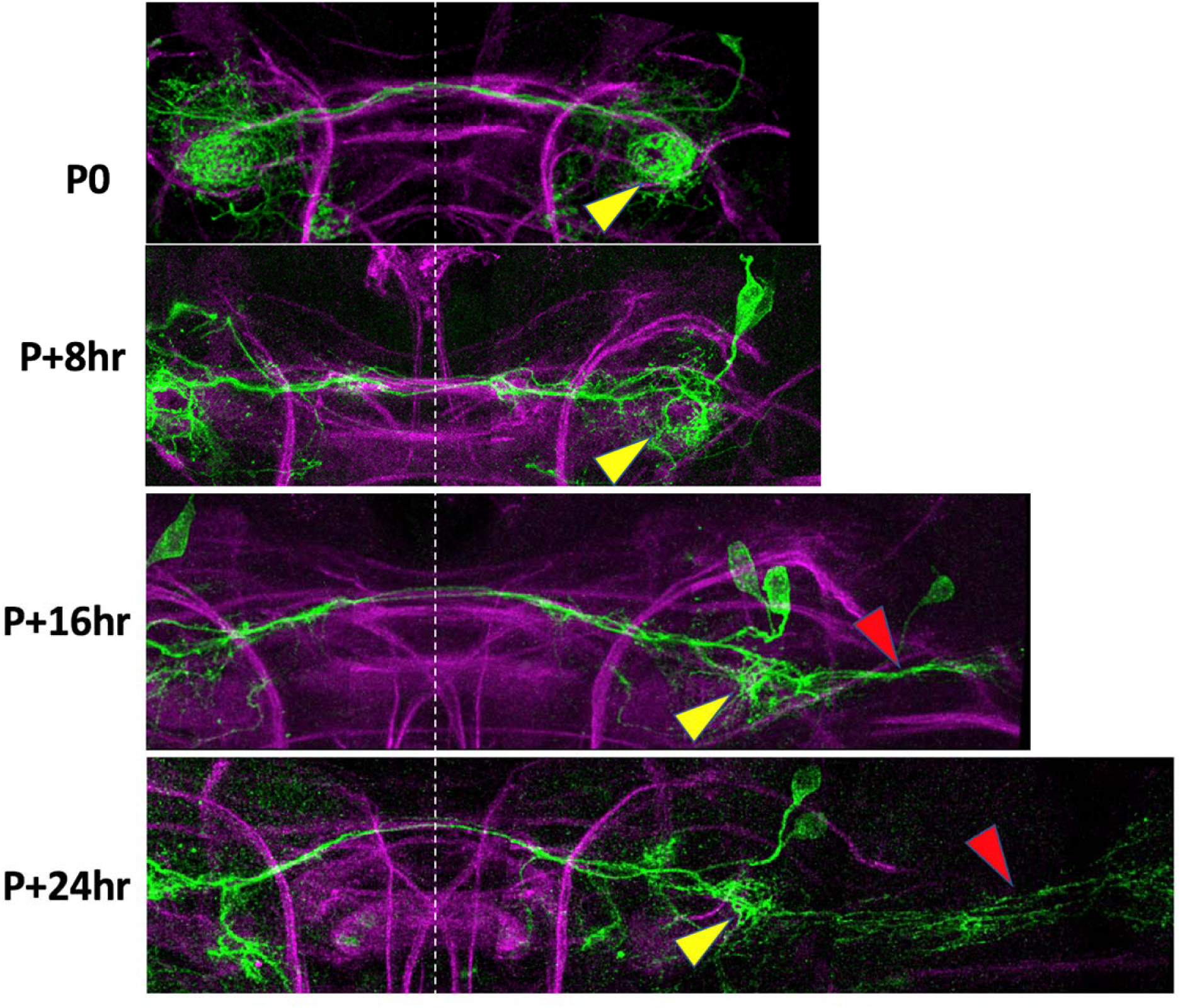
The early metamorphic development of MBIN-b1 & -b2 in hours after pupariation (P0). The yellow arrowhead marks the site where the larval cells invade the intermediate peduncle. The red arrowhead marks the outgrowth into the optic lobe. Green: Green Fluorescent Protein; magenta: fasciclin II.

**Figure 5---figure supplement 1.**
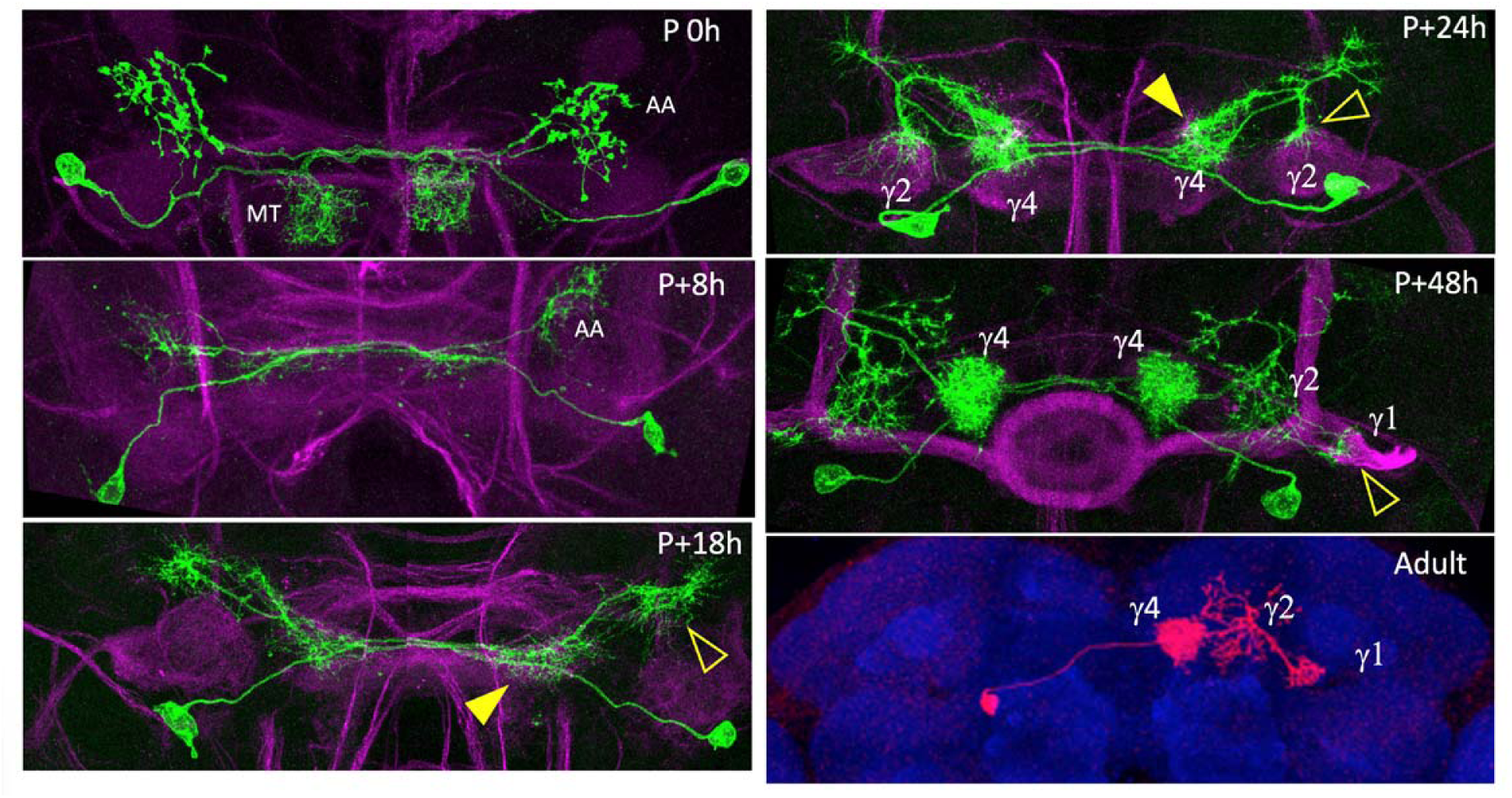
Pruning and outgrowth of MBON-j2 as it transforms into its adult form named MBON 05. At pupariation (P 0h), MBON-j2 has a dendritic arbor in the ipsilateral medial toe (MT) compartment and a contralateral axon arbor (AA). By P+8 hours, the dendritic arbor is gone and the axonal arbor has severely reduced. At P+18 hours the cell has formed contralateral outgrowth areas for new dendritic (filled arrowhead) and axonal arbors (open arrowhead). P+24 h: dendritic growth invades the γ4 compartment (filled arrowhead) while the axonal region splits into multiple growth cones, one of which invades the γ2 compartment (open arrowhead); By P + 48hr, a dendritic tuft fills the γ4 compartment and axonal arbor is in γ2, but the cell shows the delayed invasion of γ1. Adult version of the cell is an Red Fluorescent Protein version obtained by flip-switch treatment of MBON-j2 in the larva. Blue: nc82; Green: Green Fluorescent Protein; magenta: fasciclin II.

**Figure 5---figure supplement 2.**
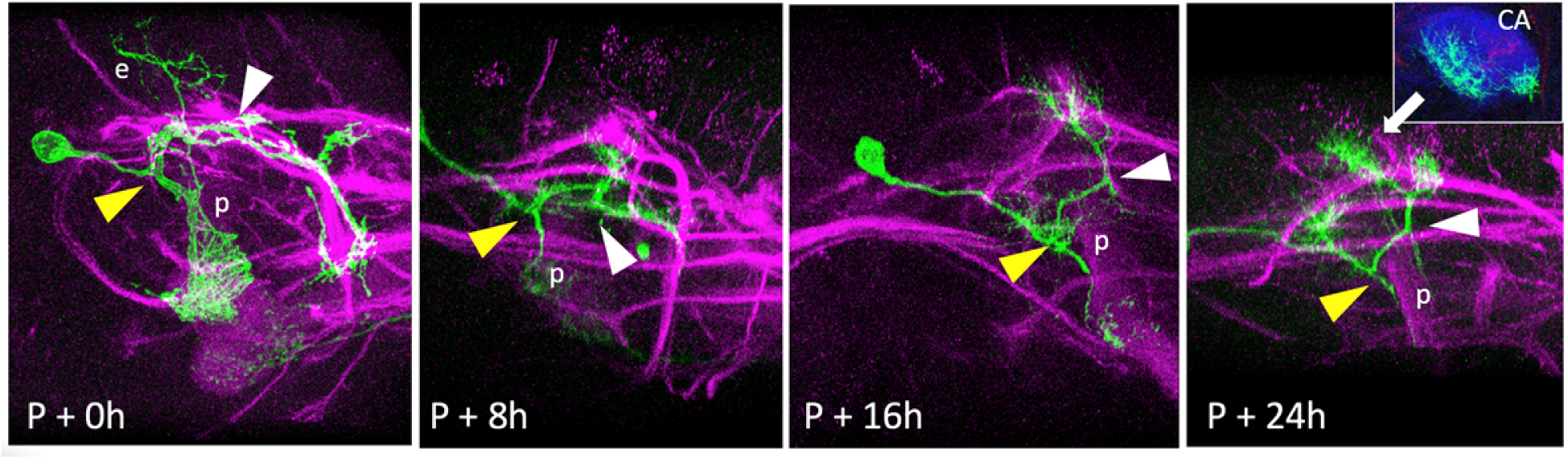
Early stages in the metamorphosis of MBON-c1. This example of MBON-c1 has an atypical ectopic branch (e) that leads to the larval calyx. Most larval cells lack this branch. Subsequent images show the progression of arbor loss and outgrowth through the 24 hours after pupariation. The yellow and white triangles show comparable junctions in the cell through time. The inset at P+24 hr is a sub-stack projection through the calyx (CA) neuropil showing growth cones invading this neuropil. p: peduncle; green, Green Fluorescent Protein; magenta: fasciculin II; blue: N-cadherin.

## Titles and Legends for Source Data Images

**Figure 2 – source data 1.** Examples of the adult anatomies of larval neurons MBIN-b1 and -b2 obtained by flip-switch mediated immortalization of expression of line SS21716 late in larval life.

**Figure 2 – source data 2.** Examples of the adult anatomies of larval neurons DAN-c1 and DAN-d1 obtained by flip-switch mediated immortalization of expression of lines MB586B and MB328B, respectively, late in larval life.

**Figure 2 – source data 3.** Examples of the adult anatomy of larval neuron OAN-e1 obtained by flip-switch mediated immortalization of expression of lines SS21716 and SS01958 late in larval life.

**Figure 2 – source data 4.** Examples of the adult anatomies of larval neurons MBIN-l1 and DAN-f1 obtained by flip-switch mediated immortalization of expression of stable spilt lines late in larval life. The anatomy of the adult form of MBIN-l1 was revealed using lines SS04484 and SS01624; that of DAN-f1 using lines MB065B and MB145B.

**Figure 2 – source data 5.** Examples of the adult anatomies of larval neurons DAN-g1 and OAN-g1 obtained by flip-switch mediated immortalization of expression of stable spilt lines late in larval life. The anatomy of the adult form of DAN-g1 was revealed using lines SS017164 and SS01755; that of OAN-g1 using lines SS20844 and SS4268.

**Figure 2 – source data 6.** Table showing the success rate for maintaining expression of the various larval neurons through metamorphosis.

**Figure 3 – source data 1.** Examples of the adult anatomy of larval neuron MBON-a1 obtained by flip-switch mediated immortalization of expression of lines SS01417 and SS00867 late in larval life. The first line also revealed an occasional adult form of MBON-a2

**Figure 3 – source data 2.** Examples of the adult anatomies of larval neurons MBON-a2 and MBON-b1,-b2 obtained by flip-switch mediated immortalization of expression of stable spilt lines late in larval life. The anatomy of the adult form of MBON-a2 was revealed using lines SS00872 and SS02006; that of MBON-b1,-b2 using lines SS01708 and SS01959

**Figure 3 – source data 3.** Examples of the adult anatomies of larval neurons MBON-d1, MBON-e2 and MBON-f2 obtained by flip-switch mediated immortalization of expression of stable spilt lines late in larval life. The anatomy of the adult form of the three neurons was revealed using lines SS01705, SS04172, and SS04328, respectively.

**Figure 3 – source data 4.** Examples of the adult anatomies of larval neurons MBON-g1 and -g2 obtained by flip-switch mediated immortalization of expression of lines SS02130 and SS02121 late in larval life.

**Figure 3 – source data 5.** Examples of the adult anatomies of larval neurons MBON-h1 and -h2 obtained by flip-switch mediated immortalization of expression of line SS01725 late in larval life.

**Figure 3 – source data 6.** Examples of the adult anatomies of larval neurons MBON-j1 and MBON-j2 obtained by flip-switch mediated immortalization of expression of lines SS01973 and SS00860 late in larval life.

**Figure 3 – source data 7.** Examples of the adult anatomies of larval neurons MBON-i1 and MBON-k1 obtained by flip-switch mediated immortalization of expression of lines SS01962 and SS04236 late in larval life.

**Figure 4 – source data 1.** Examples of the adult anatomies of larval neuron APL obtained by flip-switch mediated immortalization of expression of line SS01671 late in larval life.

## Notes

### Competing Interest Statement

The authors have declared no competing interest.

